# Computational Approach for Screening the Whole Proteome of Hantavirus and Designing a Multi-Epitope Subunit Vaccine

**DOI:** 10.1101/832980

**Authors:** Faruq Abdulla, Zulkar Nain, Md. Moyazzem Hossain, Sifat Bin Sayed, Md. Shakil Ahmed Khan, Utpal Kumar Adhikari

## Abstract

Hantaviruses are a newly zoonotic emerging group of rodent-borne viruses that have a significant impact on global public health by increasing amplitude and magnitude of outbreaks. As no permanent cure yet, it is now growing and challenging interest to develop a vaccine against Hantavirus. This study endeavored to design a robust subunit vaccine using a novel immunoinformatics approach. After meticulous evaluation, top ones from predicted CTL, HTL, and B-cell epitopes were considered as potential vaccine candidates. Among generated four vaccine models with different adjuvant, the model with TLR-4 agonist adjuvant was selected for its high antigenicity, non-allergenicity, and structural quality. The conformational B-cell epitope prediction assured its humoral response inducing ability. Thereafter, the molecular docking and dynamics simulation confirmed a good binding affinity with immune receptor TLR-4 and stability of the vaccine-receptor complex. In immune simulation, significantly high levels of IgM and IgG1 immunoglobulins, T_C_ and T_H_-cell populations, and various cytokines (i.e. IFN-γ, IL-2 etc.) are coherence with actual immune response and also showed faster antigen clearance for repeated exposures. Finally, disulfide engineering enhanced vaccine stability and *in silico* cloning confirmed the better expression in *E. coli* K12. Nonetheless, experimental validation can proof the proposed vaccine’s safety and ability to control Hantavirus infection.

## Introduction

The Hantaviruses are emerging as they cause about 200,000 outbreaks in humans annually with case fatality rates of 0.1%-50% depending on the species with the majority occurring in Asia ^1–3^; and their amplitude and magnitude of outbreaks are increasing day by day. The Hantaviruses (Genus: *Orthohantavirus*, Family: *Hantaviridae*, Order: *Bunyavirales*) are negative-stranded and trisegmented zoonotic viruses hosted by rodents, shrews, moles, bats and insects ^4,5^. Its genome composed of three molecules of negative-sense single-stranded RNA, designated S (small) encodes nucleoprotein (N), M (medium) encodes glycoproteins (Gn and Gc), L (large) encodes RdRp or L protein ^6^. The discovery of Hantaviruses in the Old and New World was issued by the two major outbreaks in the past century. Although the first clinical recognition of HFRS was in 1931 in northeast China ^7^, it became to the attention of western physicians when 3200 United Nations troops fell ill in Korea between 1951 to 1954 ^8^. Secondly in 1993, the major outbreak of previously unrecognized syndrome occurred in the Four Corners region of the United States and was referred as Four Corner disease and later it called Hantavirus cardiopulmonary syndrome (HCPS) ^9^. Therefore, Hantaviruses can cause two human acute febrile diseases so-called hemorrhagic fever with renal syndrome (HFRS) in the Old World and Hantavirus cardiopulmonary syndrome (HCPS) in the New World ^10^. The Old World and New World Hantaviruses exhibit a similar organization of nucleotide sequences and similar aspects in their lifecycle but they induce different diseases ^9^. China classified the HFRS as a class B notifiable disease ^11^ and considered it as a severe health problem ^7,12^ with a total of 112,177 cases and 1,116 deaths over the past ten years ^13,14^. From 1978 to 1995, 3,145 HFRS cases with morbidity of 1.7% were reported in Asian Russia ^15^ and in Korea, 300-500 cases are reported annually with a mean case fatality of 1% ^3^. Also, in Europe, over 3,000 HFRS cases are diagnosed annually ^3^ and over 2,800 cases were reported in Latvia ^16^. An important HFRS case has been reported in Ecuador ^17^. Furthermore, HFRS cases were reported in Vietnam, Singapore, Thailand, India, and Sri Lanka, Finland, Sweden, France, Germany, Balkans, Czech Republic, Switzerland, Poland, Greece, Lithuania, Estonia, Slovenia, Turkey, United Kingdom, and all about the African countries ^3^. In contrast, the United States listed the HCPS as a notifiable disease since 1995 ^18^ and 624 HCPS cases have been reported between the period 1993 to 2013 ^3^. In Chile, 837 HCPS cases have been recorded with a fatality rate of 36.1% during 2013 and 1,600 cases have been reported in Brazil before 2013 ^3^. Argentina reported annually 100-200 HCPS cases and some cases occurred in Canada ^3^. Furthermore, the serological evidence and cases of HCPS have been discovered in Central America, Bolivia, Colombia, French Guiana, Peru, Uruguay, Paraguay, Venezuela, and Suriname ^3,19–21^.

In 1976, Hantaan virus (HTNV) with its reservoir, striped field mouse (*Apodemusagrarius*) of *Muridae* family were reported by Lee *et* al. as the first etiological agent of HFRS in South Korea along the Hantaan River ^22^ and later in China and Russia ^3^. In 1930s, the milder form of HFRS called Nephropathia Epidemica (NE) was first described in Sweden and thousands of infection cases occur annually throughout Europe ^23^ and it is an etiological agent Puumala virus (PUUV) was found in bank voles (*Myodesglareolus*) in Finland in 1980 ^24^. The Puumala virus (PUUV) is also an agent of HFRS and recently HCPS was reportedly induced by PUUV in Germany ^25^. The Seoul virus (SEOV) hosted by rates is the second most significant pathogenic agent of HFRS found predominantly in Korea and worldwide ^26,27^. Further, the Dobrava-Belgrade virus (DOBV) ^28^ and Tula virus (TULV) were found as human pathogenic agents of HFRS in Europe ^29^. The causative agent Thailand virus (THAIV) and THAIV-like virus Anjozorobe virus (ANJOV) were in Thailand and Madagascar, respectively ^6,30–32^. Heinemann *et* al. reported the Bowe virus (BOWV) as HFRS related human pathogenic agent ^2^ and also the Sangassou virus (SANGV) have described as HFRS agents and found in West Africa (Guinea) ^33^. In contrast, the Sin Nombre virus (SNV) and Andes virus (ANDV) were discovered as etiological agents of HCPS in North and South America, respectively ^3,34^ where only ANDV has the characteristics of person-to-person transmission. Subsequently, there are about 43 strains have been identified in the Americas and 20 strains of them are associated with human disease ^3^. In Brazil, the Araraquara virus (ARAV) is a top virulent agent of HCPS with a case fatality rate of 50% ^20^. Furthermore, HCPS was also reported for the cause of Bayou virus (BAYV), Black Creek Canal virus (BCCV), Choclo virus (CHOV), Bermejo virus (BMJV), Lechiguanas virus (LECV), Oran virus (ORNV), Maciel virus (MCLV), Laguna Negra virus (LNV), Hu39694 virus, Neembucu virus, Cano Delgadito virus (CADV), Tunari virus, Blue River virus, and El Moro Canyon virus (ELMCV) ^3,35–38^.

The main route of the human infection by Hantaviruses is the inhalation of aerosols contaminated with the virus concealed in the excreta, saliva, and urine of the infected animals ^10^. The human kidney and lung are the main targeted organs of HFRS-associated and HCPS-associated viruses, respectively ^3^. In humans, Hantaviruses mainly infect vascular endothelial cells and dysfunctioning them in capillaries and small vessels. Therefore, the dramatic increase in vascular permeability is the basic pathology of hantavirus-associated diseases ^12^. Basically, HFRS patients have manifested five clinical phases including fever, hypotensive shock, oliguric, polyuric, and convalescent and on the other hand, prodromal, cardiopulmonary, and convalescent are the three clinical phases of HCPS patients ^3^. The incidence in males is over three times greater than that in females ^3^. Acute encephalomyelitis, bleeding, multiorgan dysfunction, pituitary hemorrhage, glomerulonephritis, pulmonary edema, shock, respiratory distress syndrome, disseminated intravascular coagulation, and lethal outcome are the main complications of HFRS and in contrast, renal insufficiency, thrombocytopenia, bleeding, myalgia, headache, nausea, vomiting, diarrhea, shock, and lethal outcome are associated with HCPS ^3^. HCPS is a fast-evolving disease with a high case fatality rate and a patient can evolve from acute febrile illness to severe pneumonia with respiratory failure and cardiogenic shock ^9^. However, against this severely emerging virus, there are only some inactivated vaccines have been developed from Hantavirus in cell cultures or the rodent brain, and a few of these have been licensed for humans in Korea and China ^35^.

The immune response plays an important role in the pathogenesis of Hantavirus infection ^3^ and conversely plays a critical role in fighting tumors and viral infections ^39^. In modern times, immunotherapy is a powerful and efficient strategy for the prevention of infectious diseases. Recently, the multi-epitope vaccine is an ideal approach for the prevention and treatment of tumors or viral infections ^40–44^. There are many multi-epitope vaccine design studies involving various viruses like HIV ^45^, Dengue virus ^46^, Hepatitis B virus ^47^, Hepatitis C virus ^48,49^, Ebola virus ^50^, Chikungunya virus ^51^, Avian influenza A (H7N9) virus ^52^, Zika virus ^53^, Classical swine fever virus ^54^, Nipah virus ^55^, and Norovirus ^56^. An ideal multi-epitope vaccine should be designed to include a series of or overlapping epitopes so that its every basic unit called antigenic peptide fragment can elicit either a cellular or a humoral immune response against the targeted tumor or virus ^39^. In present, the multi-epitope vaccine approach has fascinated more global attention over traditional or single-epitope vaccine as it has a unique design mechanism with some properties ^39^: (a) consist of multiple MHC-restricted that can be recognized by TCRs of multiple clones from various T-cell subsets; (b) consist of T_C_, T_H_, and B-cell epitopes that can induce strong cellular and humoral immune response simultaneously; (c) consist of multiple epitopes from different tumor/virus antigens that can expand the spectra of targeted tumors or viruses; (d) introduce some components with adjuvant capacity that can enhance the immunogenicity and long-lasting immune responses; and (e) reduce unwanted components that can trigger either pathological immune responses or adverse effects. A well-designed multi-epitope vaccine should become a powerful prophylactic and therapeutic agent against the targeted tumor or virus. In this study, a series of immunoinformatics approaches have been applied on the whole proteome of Hantavirus for developing a multi-epitope vaccine using the screened robust T_C_, T_H_, and B-cell vaccine candidates. Furthermore, the developed vaccine was studied for its antigenicity and allergenicity and subsequently the structural prediction was done. Moreover, the codon adaptation and *in silico* cloning, disulfide engineering, binding affinity with the immune receptor, and the molecular dynamics simulation of the best vaccine-receptor docked complex were sequentially studied. The *in silico* immune simulation has also been performed.

## Results and discussion

### Representative Proteins Selection of Orthohantavirus

In order to design a candidate vaccine, the total of 2487 protein sequences of 112 organisms of orthohantavirus was retrieved from the NIAID Virus Pathogen Database and Analysis Resource (ViPR) in a single FASTA file; where 68 organisms with 2162 proteins are classified and the remaining 44 organisms with 325 proteins are unclassified. The data overview is illustrated in Fig. 1 after rearranging the retrieved sequences according to their category, organism, and epidemic nature.

**Figure 1:**
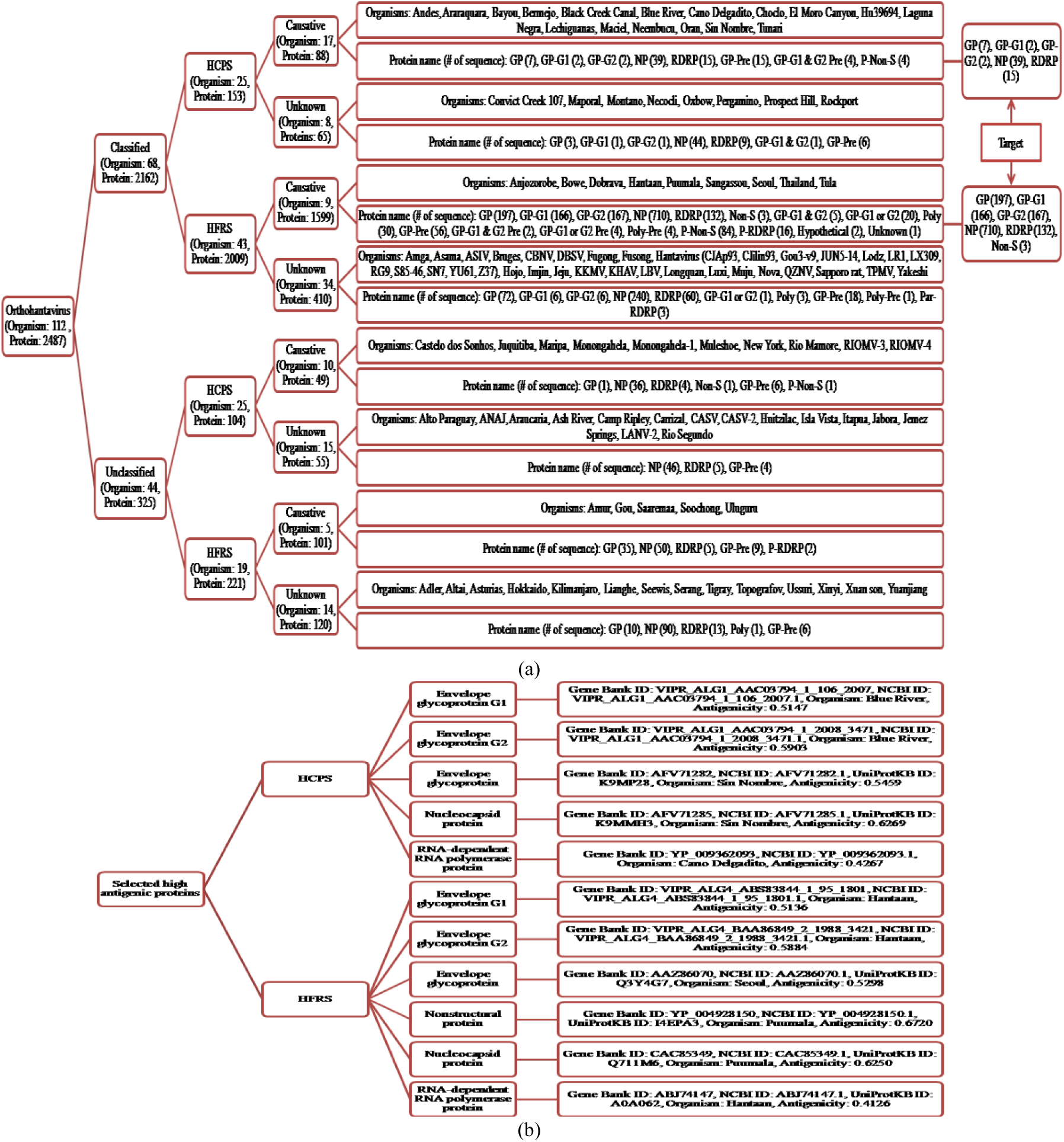
Overview of the analyzed data where (a) indicates the classification of the whole proteome of the Hantavirus retrieved from ViPR database and (b) represents the summary of the highest antigenic proteins.

In Fig. 1(a), only 27 organisms (classified: 17, unclassified: 10) with 137 proteins (classified: 88, unclassified: 49) are causative for human HCPS and 23 organisms (classified: 8, unclassified: 15) with 120 proteins (classified: 65, unclassified: 55) are unknown for human infection but associated with HCPS. As opposed to, only 14 organisms (classified: 9, unclassified: 5) with 1700 proteins (classified: 1599, unclassified: 101) are causative for human HFRS and 48 organisms (classified: 34, unclassified: 14) with 530 proteins (classified: 410, unclassified: 120) are unknown for human infection but associated with HFRS. Among 18 protein categories, only the envelope glycoprotein, envelope glycoprotein G1, envelope glycoprotein G2, nucleocapsid protein, RNA dependent RNA polymerase (RdRp) protein, and nonstructural protein from every group are the central target of this study. The mentioned proteins of the classified causative groups were tested for antigenicity and the 11 highest antigenic proteins were selected as an input dataset of the immunoinformatics study. The summary of the selected highest antigenic proteins is shown in Fig. 1(b).

### Prediction of Cytotoxic and Helper T-Lymphocyte Vaccine Candidates with Their Associated MHC HLA Alleles

The highest antigenic proteins were subjected to the NetCTL v1.2 server and a total of 3769 epitopes with a combined score≥0.5 were predicted. The predicted epitopes were considered to investigate their antigenicity, immunogenicity, conservancy, toxicity, and their respective MHC HLA alleles. Thereafter, the predicted epitopes were filtered according to the antigenicity≥0.4, immunogenicity>0, toxicity<0, and allele available in PDB>0 and found that 654 epitopes successfully satisfied all the criteria. Later, these 654 epitopes were tested their allergenicity and found that only 380 epitopes showed a non-allergic nature. Finally, among those 380 epitopes, the best one epitope from each protein was selected as a vaccine candidate that has characteristics better than others of that protein (Tab. 1). Again, the highest antigenic proteins were submitted to the IEDB MHC II binding tool and a total of 2197 epitopes were predicted with consensus percentile rank≥2. The predicted epitopes were tested their antigenicity, conservancy, and toxicity and found that 1245 epitopes were satisfied antigenicity≥0.4 and non-toxin criteria. Afterword, these 1245 epitopes were evaluated for their allergenicity and observed that 771 epitopes were found to be non-allergic. Later, these 151 epitopes were investigated for their IL-10 and IFN-γ inducing criteria. Finally, the best one epitope from each protein was selected as vaccine candidate that satisfied IL-10 and IFN-γ inducing criteria and have other characteristics better than the remaining epitopes of that protein (Tab. 2).

**Table 1:**
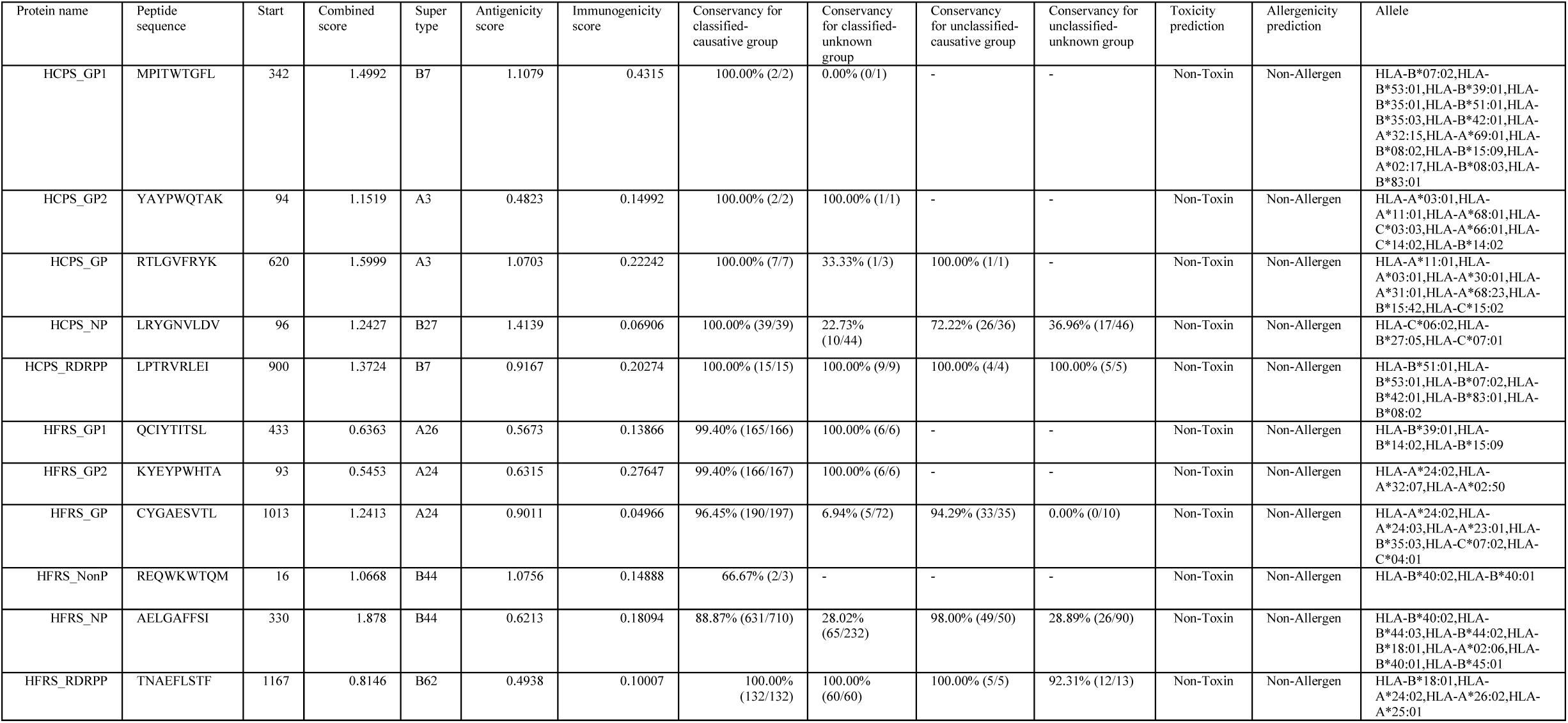
Potential CD8+ T-cell vaccine candidates and their characteristics corresponding to each protein of orthohantavirus.

**Table 2:**
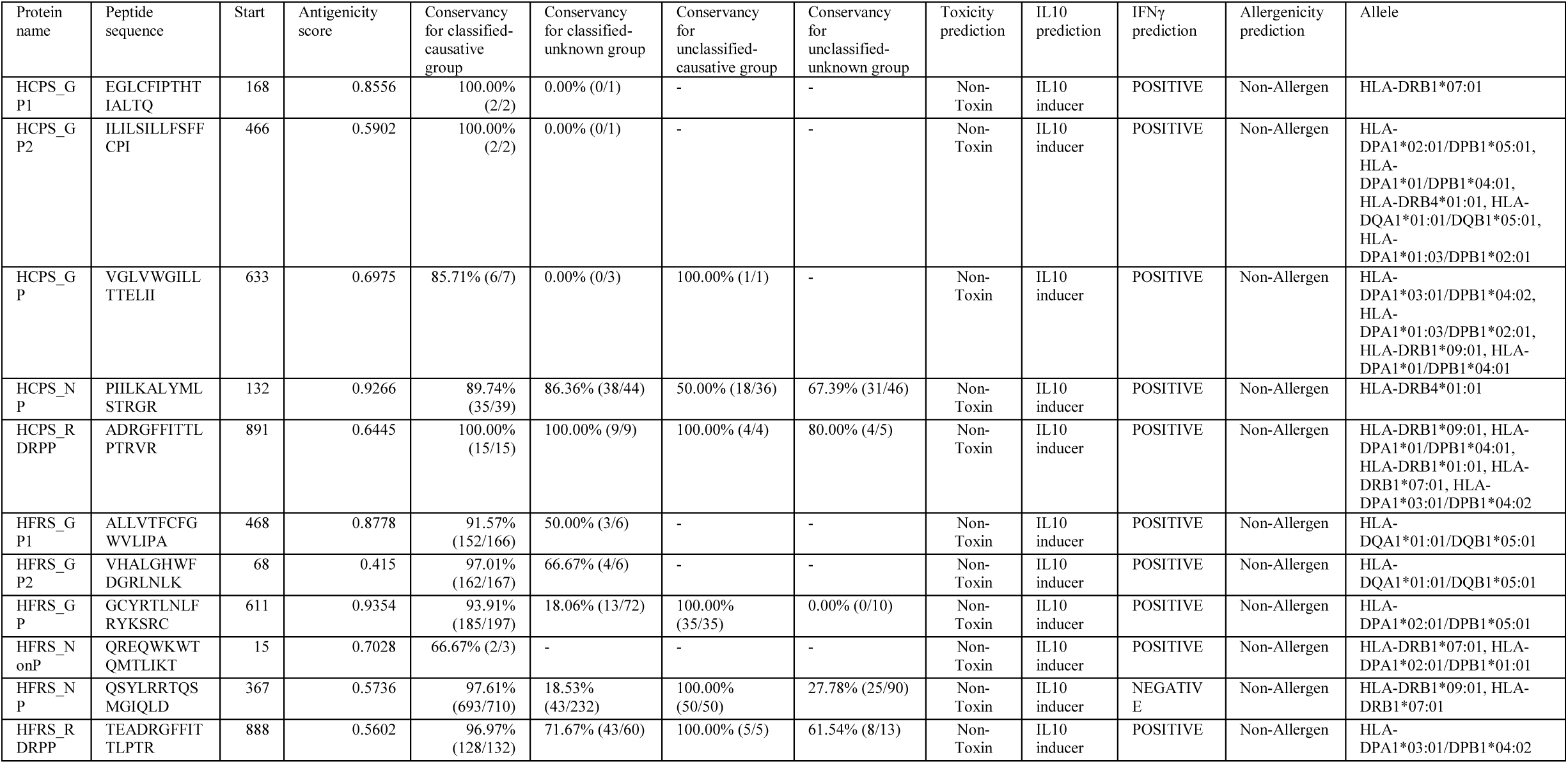
Potential CD4+ T-cell vaccine candidates and their characteristics corresponding to each protein of orthohantavirus.

### Prediction of Linear B-Lymphocyte Vaccine Candidates

The selected highest antigenic 11 proteins were submitted to the LBtope server and a total of 2939 epitopes were predicted. The predicted epitopes were considered for the evaluation of their antigenicity, conservancy, and toxicity. According to the antigenicity≥0.4 and non-toxin criteria, only 1789 epitopes have remained and were taken into account for investigating their allergenicity and found that 1011 epitopes have non-allergic nature. Subsequently, among the non-allergic epitopes, from each protein the best one epitope was selected as a vaccine candidate that significantly satisfies all the characteristics than other epitopes of that protein (Tab. 3).

**Table 3:**
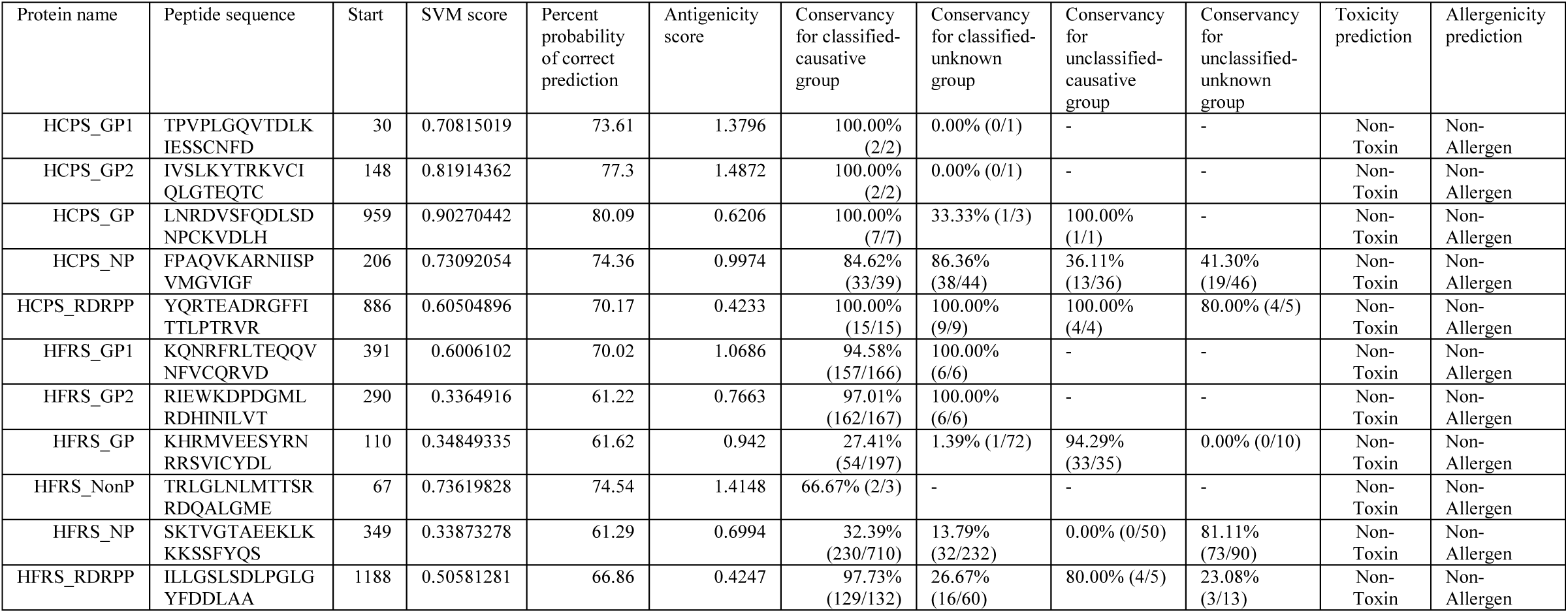
Potential B-cell vaccine candidates and their characteristics corresponding to each protein of orthohantavirus.

### Population Distribution Analysis

A combined population distribution analysis was performed for the finally selected CTL & HTL vaccine candidates and their respective HLA alleles. The analysis result showed excellent population coverage in different epidemic and non-epidemic countries, areas, and ethnic groups (Tab. S1). Among epidemic regions, the result exhibited the maximum coverage of the population in Finland (99.62%), closely followed by Germany (99.45%), Poland (99.31%), Czech Republic (99.27%), Mexico (99.19%), Sweden (99.14%), Russia (98.98%), Japan (98.91%), France (98.86%), South Korea (98.50%), United States (98.45%), and others (Fig. 2(a)). Herein, we mentioned 16 different geographical areas capturing the world where the maximum percentage of cumulative population coverage in Europe (99.10%), nearly followed by East Asia (98.72%), North America (98.43%), and others (Fig. 2(b)). The 97.94% population of the world covered by predicted vaccine candidates (Fig. 2(b)). The population coverage analysis revealed that the MHC HLA alleles corresponding to predicted T-cell vaccine candidates are well & widely distributed throughout the world, which is a robust property of rational vaccine design ^57,58^.

**Figure 2:**
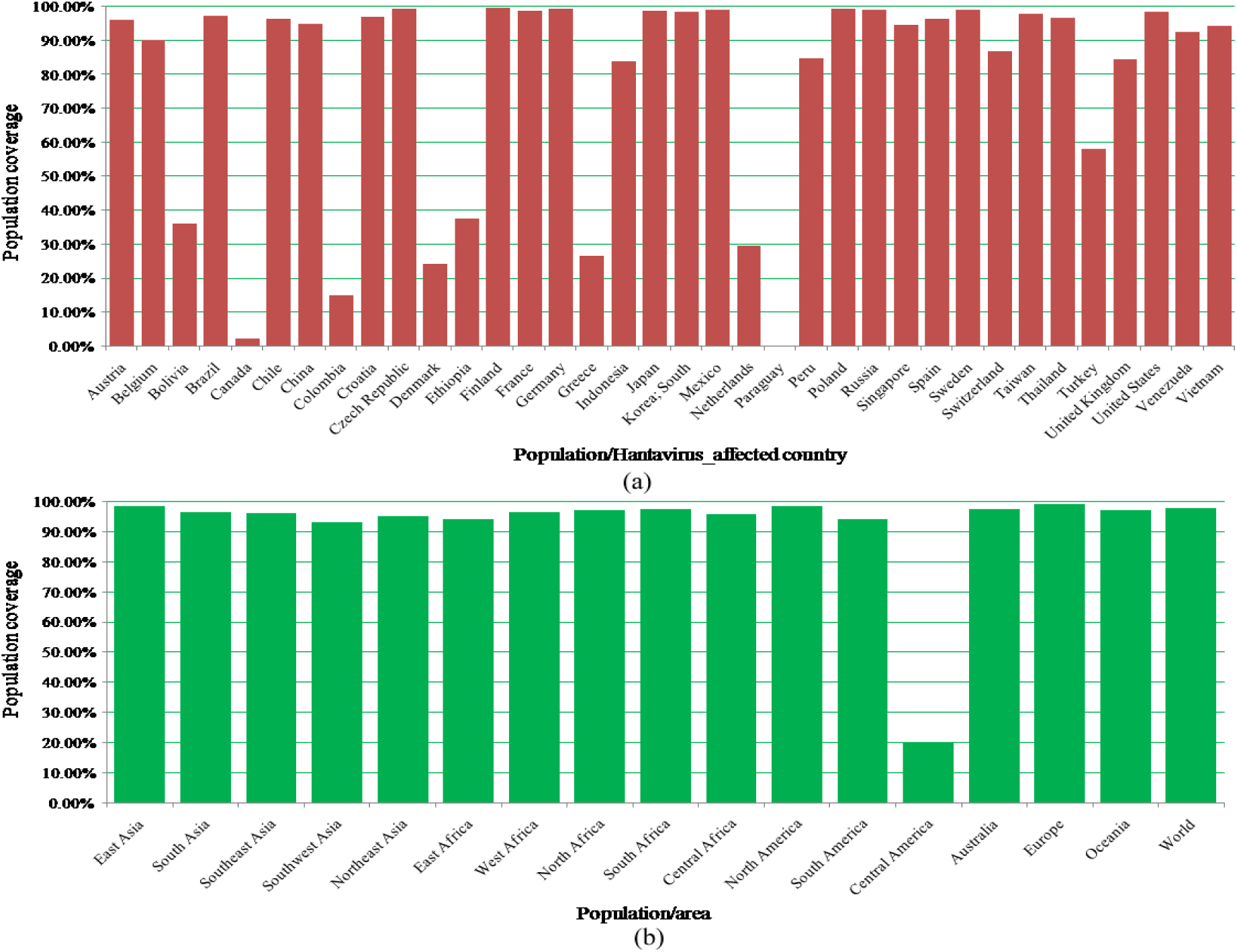
Percentage of population coverage. (a) Percentage of population coverage in the Hantavirus affected countries. (b) Percentage of population coverage in 16 different areas covering the world as well as in the whole world.

### Molecular Docking Analysis of MHC HLA Alleles-Peptides Interactions

The binding ability of the predicted vaccine candidates with their associated MHC HLA alleles should be checked for their robustness. To do this, the predicted vaccine candidates were modeled through PEPFOLD v3.5 and their associated lowest percentile ranked allele proteins were downloaded from PDB. At first, the control score of the allele proteins with their experimental ligand was examined. Thereafter, the molecular dockings of the peptide-allele complexes were done through AutoDock & AutoDock Vina and the docking scores were compared with their control scores (Fig. 3). The vaccine candidates exhibited good binding energy with their respective HLA-alleles. The epitope-allele docked complexes were shown in Fig. 4.

**Figure 3:**
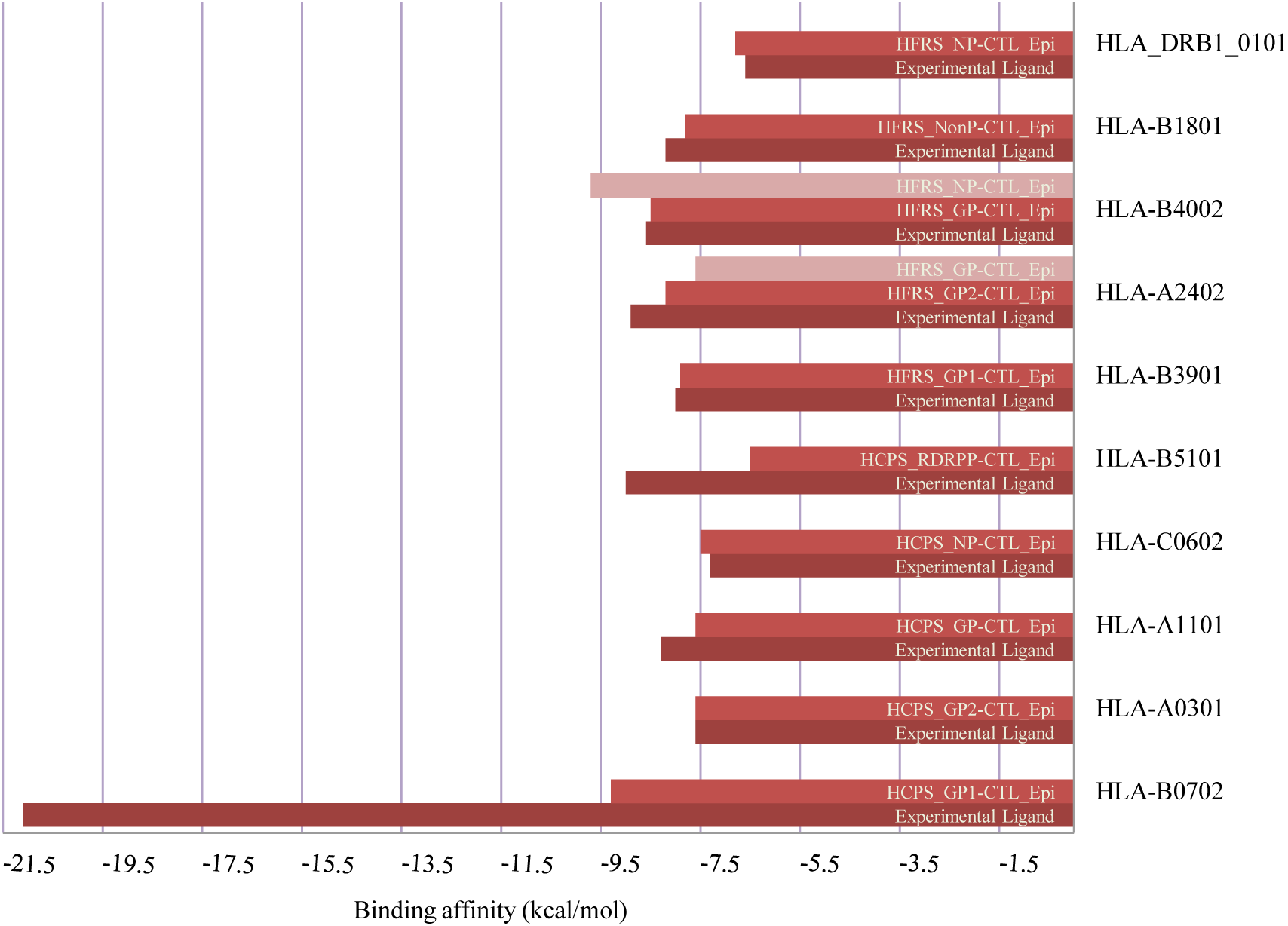
Epitope-allele docking scores comparing with the experimental peptide-allele docking scores (dark red color indicating the epitope-allele docking score and light red color indicating the experimental peptide-allele docking score).

**Figure 4:**
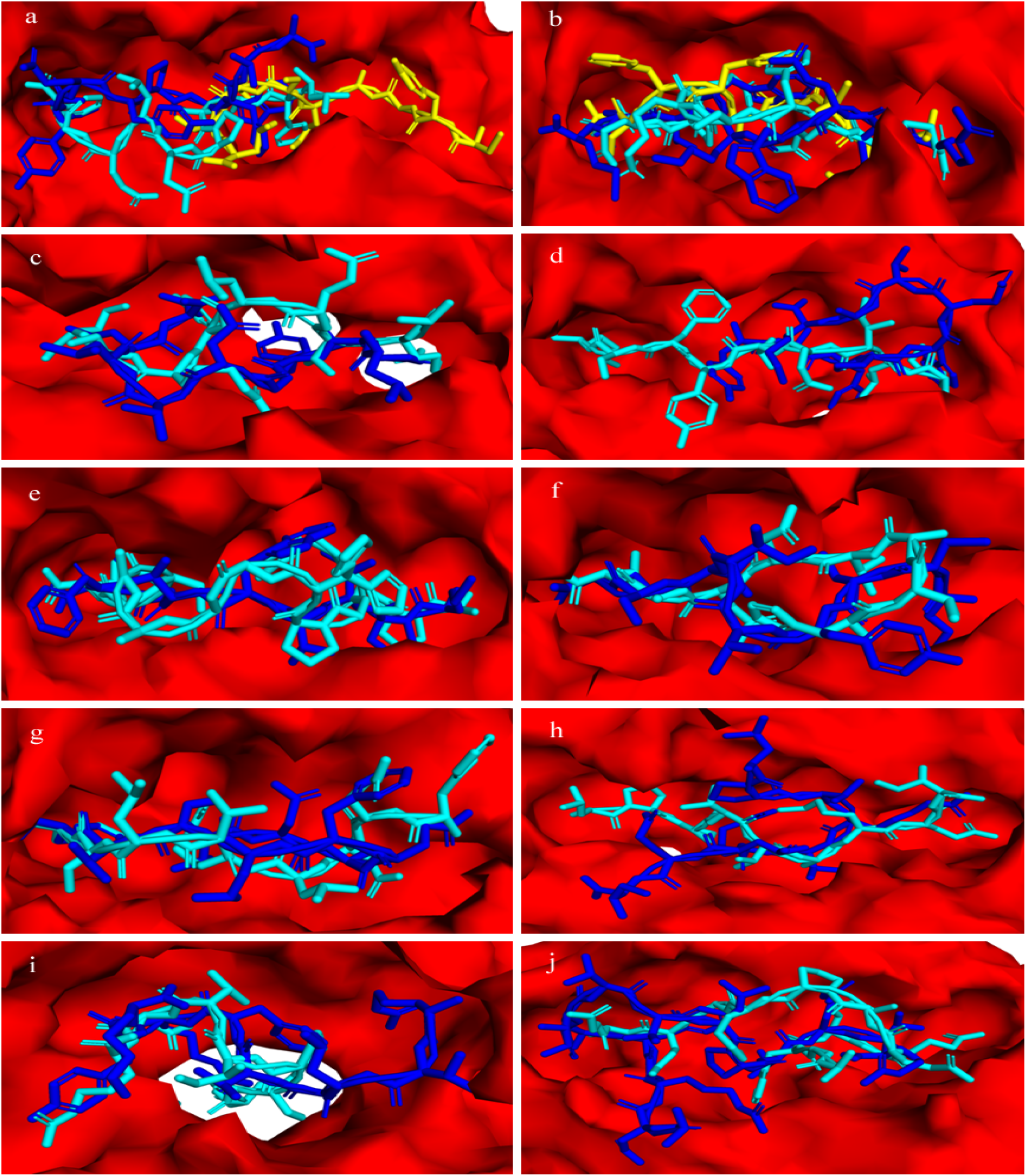
Epitope-allele docked complexes. In all the figures, red color represents the allele protein, cyan color indicates the experimental ligand, and blue & yellow colors highlight the epitopes. The figure (a) shows the docked complexes of HLA-A2402 with HFRS_GP2-CTL_Epitope (blue) and HFRS_GP-CTL_Epitope (yellow) and the figure (b) shows the docked complexes of HLA-B4002 with HFRS_GP-CTL_Epitope (blue) and HFRS_NP-CTL_Epitope (yellow). The figures (c-i) highlight the docked complexes of HLA-C0602, HLA-A0301, HLA-B0702, HLA-B3901, HLA-B1801, HLA-B5101, and HLA-A1101 with HCPS_NP-CTL_Epitope, HCPS_GP2-CTL_Epitope, HCPS_GP1-CTL_Epitope, HFRS_GP1-CTL_Epitope, HFRS_NonP-CTL_Epitope, HCPS_RDRPP-CTL_Epitope, and HCPS_GP-CTL_Epitope, respectively. The figure (j) shows the docked complex between HLA_DRB1_0101 and HFRS_NP-CTL_Epitope.

### Cluster Analysis of the MHC Restricted HLA Alleles

The MHCcluster v2.0 generated a correlation heat-map for clustering both the MHC-I and MHC-II HLA molecules interacted with the predicted robust vaccine candidates. The generated heat-map is illustrated in Fig. 5 showing a satisfactory functional relationship among detected allele molecules as the red-colored zones indicating strong correlation and gradually yellow-colored zones indicating weaker correlation ^59^.

**Figure 5:**
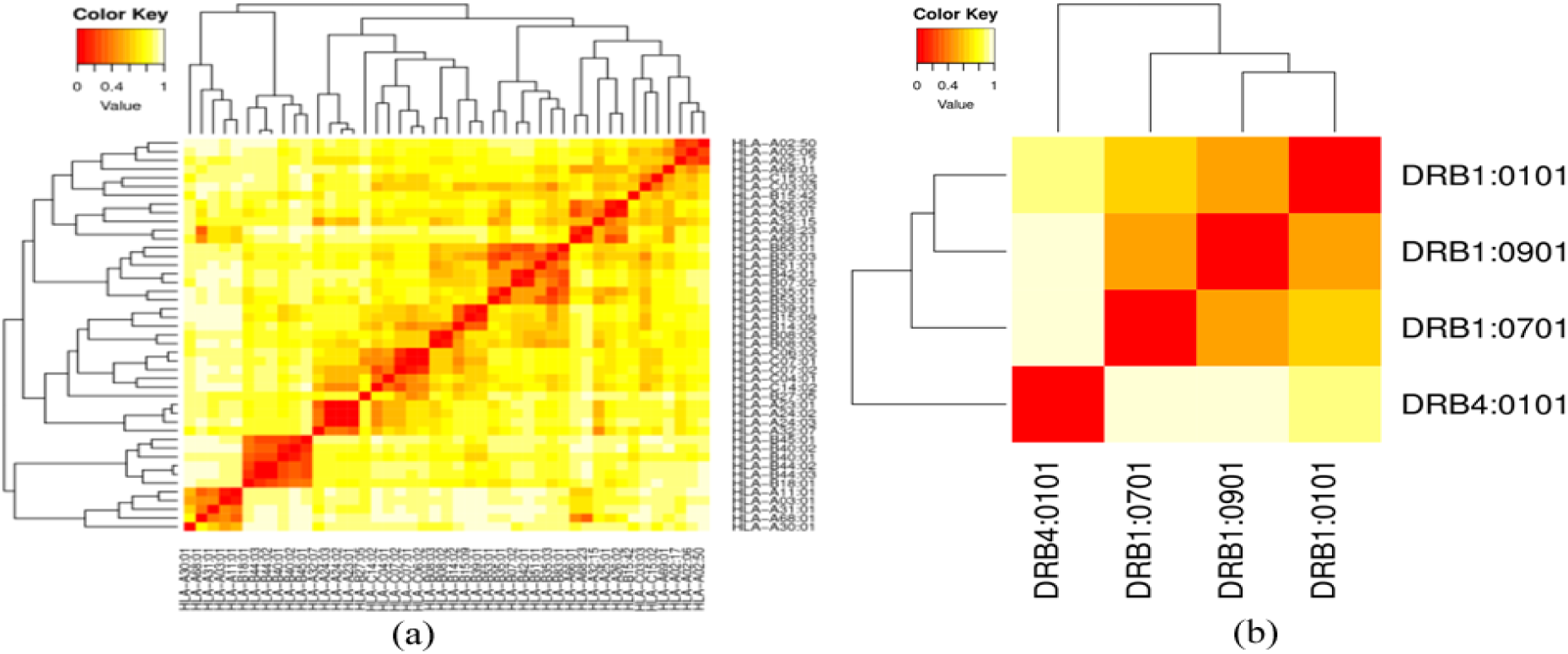
Cluster analysis of the MHC HLA alleles where (a) is the clustering of MHC-I HLA alleles and (b) is the clustering of the MHC-II HLA alleles (red colors is the indication of strong interaction, while the yellow zone indicating the weaker interaction).

### Multi-Epitope Subunit Vaccine Protein Construction

The immune response plays a critical role in fighting viral infections and its molecular and cellular mechanisms induced by the multi-epitope vaccine ^39^. Therefore, extracted robust vaccine candidates of total 11 CTL, 11 HTL, and 11 BL epitopes were sequentially fused together with the help of suitable linkers for the long chain of multi-epitope vaccine protein. The linkers GGGS, GPGPG, and KK were used to connect the intra-CTL, intra-HTL, and intra-BL epitopes, respectively. A multi-epitope subunit vaccine must contain a strong immunostimulatory adjuvant for enhancing the immunogenicity and activating long-lasting innate and adaptive immune response ^39,60^. Four vaccine models of length 654, 692, 686, and 777 amino acids were designed by adding TLR4 agonist, β3-defensin, β3-defensin with RR residues, and 50S ribosomal protein L7/L12 at the N-terminal site of the vaccine protein using EAAAK linker, respectively. The lengths of the designed vaccine models are not too long comparing to the other developed multi-epitope vaccine against *Fasciola gigantic, Schistosoma mansoni* and *Onchocerca volvulus* of length 765, 617 and 599, respectively ^61–63^. Shanmugam *et al.* reported that synthetic TLR4 can be used as a novel adjuvant ^64^. The β-defensin can recruit the Naïve T-cell and immature dendritic cells (DC) at the site of infection through the CCR6 receptor and it can also provide an adaptive immune response and innate host response in microbial infection ^65^. The 50S ribosomal protein L7/L12 act as an immunoadjuvant for dendritic-cell (DC) based immunotherapy and it is capable of inducing DC maturation ^66^. The schematic diagram of the vaccine construct is represented in Fig. 6.

**Figure 6:**
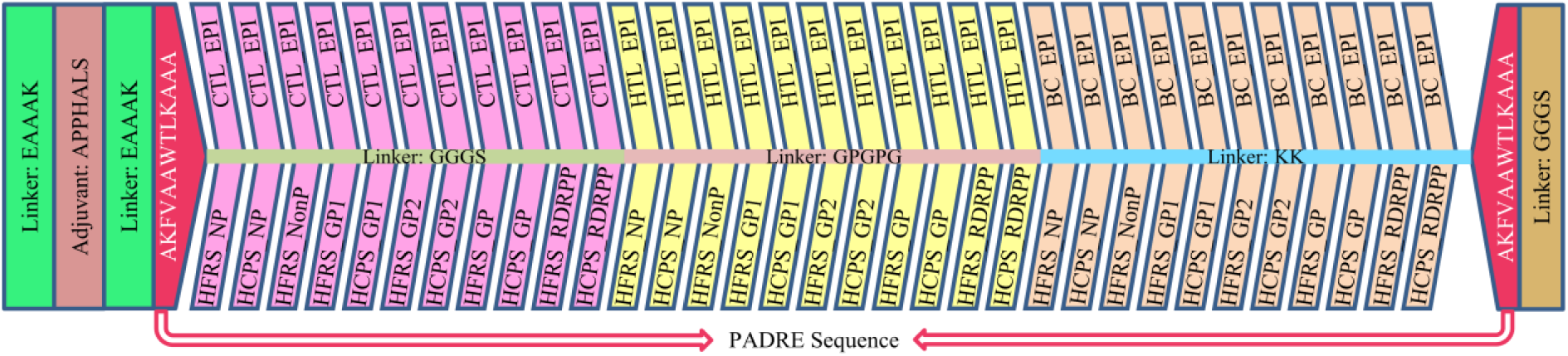
Schematic diagram of the multi-epitope subunit vaccine construct. An adjuvant was added at the N-terminal site of the construct with the help of EAAAK linker and the T_C_, T_H_, and BL epitopes were merged with the help of GGGS, GPGPG, and KK linkers, respectively.

### Immunological Comparison of Vaccine Models

Immunological properties such as antigenicity and allergenicity were evaluated for all vaccine models and the results were illustrated in Fig. 7(a). The high antigenicity indicates the better ability to induce an immune response. As the allergy involves a series of complex reactions by the immune system due to the foreign antigenic substance and results in sneezing, wheezing, skin rash, and swelling of the mucous membrane, the non-allergic vaccine is safe for human life ^67^. All the vaccine models showed antigenic and non-allergic in nature. The vaccine model-1 was characterized by higher antigenicity of 0.9876 but the higher non-allergen score of −0.917 was assigned to the vaccine model-3.

**Figure 7:**
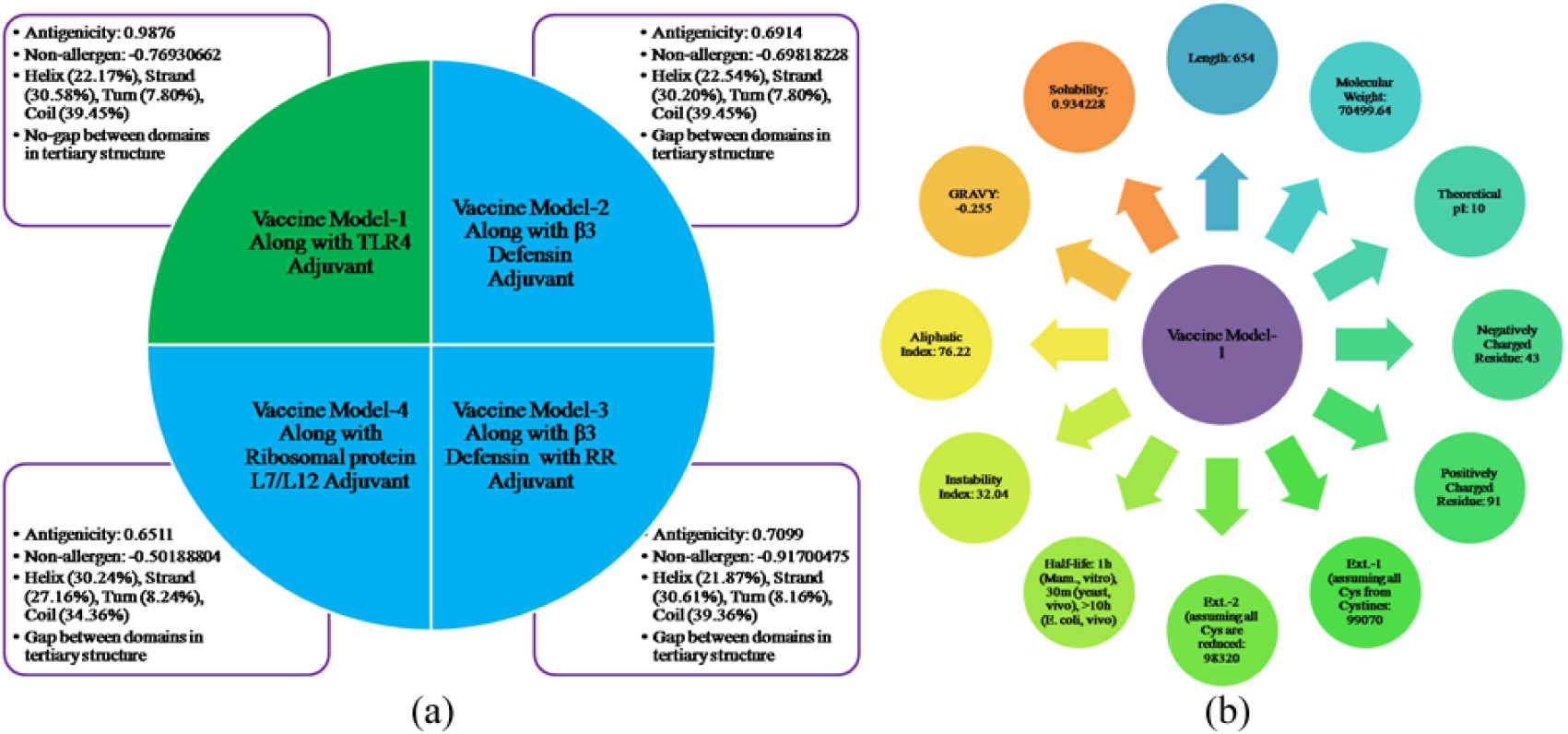
Immunological and tertiary structural properties of the vaccine constructs. (a) Comparison of the vaccine models by immunological and structural properties. (b) Physicochemical properties of the finally selected vaccine model-1.

### Structural Comparison of Vaccine Models

The secondary structural features of the vaccine models were assessed from SOPMA server using their amino acid sequences. The predicted results were illustrated in Fig. 7(a). Thereafter, the vaccine models were submitted to RaptorX server for predicting their 3D structures. The predicted results were checked for the gap between domains of the predicted 3D models and observed that only the vaccine model-1 exhibited no gap between domains (Fig. 7(a)). The 3D protein models having a gap between domains create problems in the refining stage of those models ^68^. Therefore, the vaccine model-1 was selected as the final vaccine construct for further immunoinformatics approaches. A probability score graph of occurrence of helix, strand, turn, and coil at each amino acid position in the secondary structure of the final vaccine construct is shown in Fig. 8(a). The tertiary structure of the final vaccine construct was predicted as 2 domains (5j81:A, 4ind:A) and modeled using 4ind:A as the best template with a score of 50. All of the 654 amino acids were modeled, but 10% of residues were predicted as disordered. The p-value and uGDT value are the quality measures in homology modeling, lower p-value and higher uGDT value confirm the quality of the modeled structure ^69^. The p-value and uGDT values obtained for the modeled structure were 8.84×10^−5^ and 120, respectively, which are sufficiently low and large, respectively.

**Figure 8:**
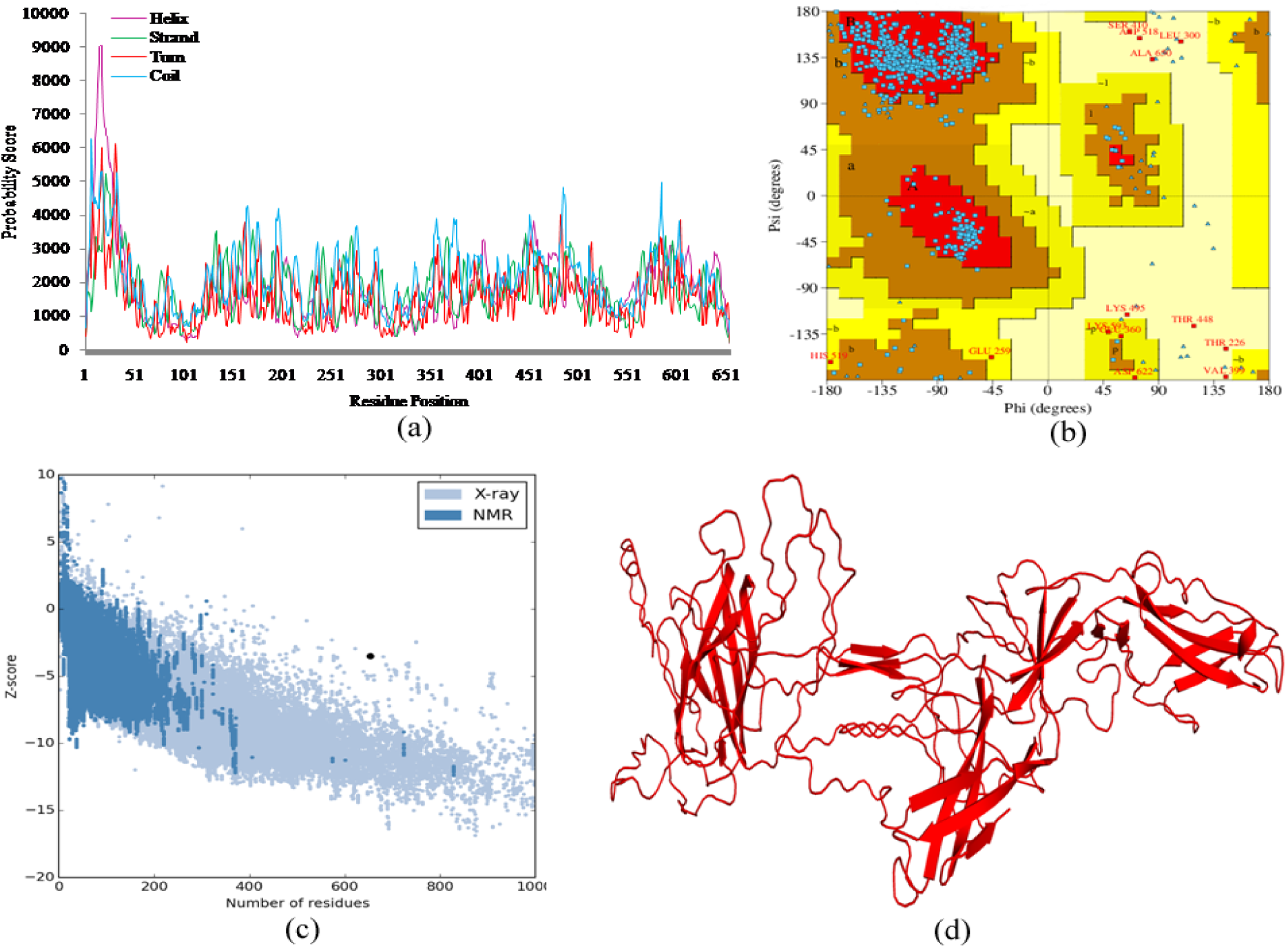
Structural assessment of the final vaccine construct. (a) Probability score graph of occurrence of helix (purple), strand (green), turn (red), and coil (light blue) at each amino acid position in the secondary structure of the final vaccine construct. (b) Ramachandran plot of the refined model showing 86.2%, 11.3%, 1.0%, and 1.3% residues were found in most Rama-favored regions, additional allowed regions, generously allowed regions, and disallowed regions respectively. (c) The overall quality Z-score plot from ProSa server showing Z-score of −3.51. (d) Refined 3D structure of the final vaccine construct.

### Refinement and Quality Assessment of the Tertiary Structure of the Final Vaccine Construct

Structural refinement with GalaxyRefine can improve the structural quality by increasing the number of residues in the favored region. GalaxyRefine provided five models with various quality assessment parameters (Tab. S2). Among them, the model 4 was taken into consideration as the best 3D structure of the final vaccine construct for further analysis according to various quality parameters including GDT-HA (0.9209), RMSD (0.502), MolProbity (2.432), Clash score (28.6), Poor rotamers (0.8), and Rama favored (91.9). For the structural validation of the refined model, the Ramachandran plot analysis was done and found that 86.2%, 11.3%, 1.0%, and 1.3% residues in Rama-favored regions, additional allowed regions, generously allowed regions, and disallowed regions, respectively (Fig. 8(b)). The overall quality and potential errors in a crude 3D model were verified through ERRAT and ProSA-web server. The overall quality factor of the refined model was 52.13% using ERRAT. While ProSA-web has shown the Z-score of −3.51 of the refined model which is out of range that commonly found in the case of native proteins for comparable size but it is supported by experimentally validated structures and close to the database average (Fig. 8(c)) ^70^. The refined 3D model is illustrated in Fig. 8(d).

### Assessment of Primary Sequence Features of the Final Vaccine Construct

The various primary sequence features of the final vaccine construct were evaluated through ExPASy ProtParam server (Fig. 7(b)). The molecular weight of the final vaccine construct was found to be 70.5 kDa which is the indication of good antigenic nature of the final vaccine construct ^70^. The proteins having <110 kD molecular weight are believed to be good vaccine candidates ^71^. The theoretical isoelectric point (pI) is valuable to provide a buffer system for vaccine purification which was computed to be 10 showing slightly basic in nature of the construct. Moreover, the estimated half-life was found to be 1 hour in mammalian reticulocytes, *in vitro*; while 30 minutes in yeast and >10 hours in *E. coli, in vivo*. The instability index was assessed to be 32.04 which represents the stable nature of the final vaccine construct ^72^. The score of the aliphatic index was 76.22 which is the indication of the thermostable nature of the final vaccine construct; the higher aliphatic index is associated with more thermostability ^60^. The estimated grand average of hydropathicity (GRAVY) value was calculated as −0.255, negative GRAVY value declared the final vaccine construct is hydrophilic in nature and it has good interaction with a water molecule ^60^.

### Mapping of Conformational B-cell Epitopes in Final Vaccine Construct

B-cells are the key player of humoral immunity and epitope corresponding to the B-cell receptor plays an important role in vaccine design following antibody production ^60^. The discontinuous B-cell epitopes were searched in the final vaccine construct using the IEDB ElliPro tool and found 10 significant discontinuous B-cell epitopes with length range 4-110 and score range 0.557-0.811 (Tab. S3). The structural view of those epitopes was illustrated in Fig. 9.

**Figure 9:**
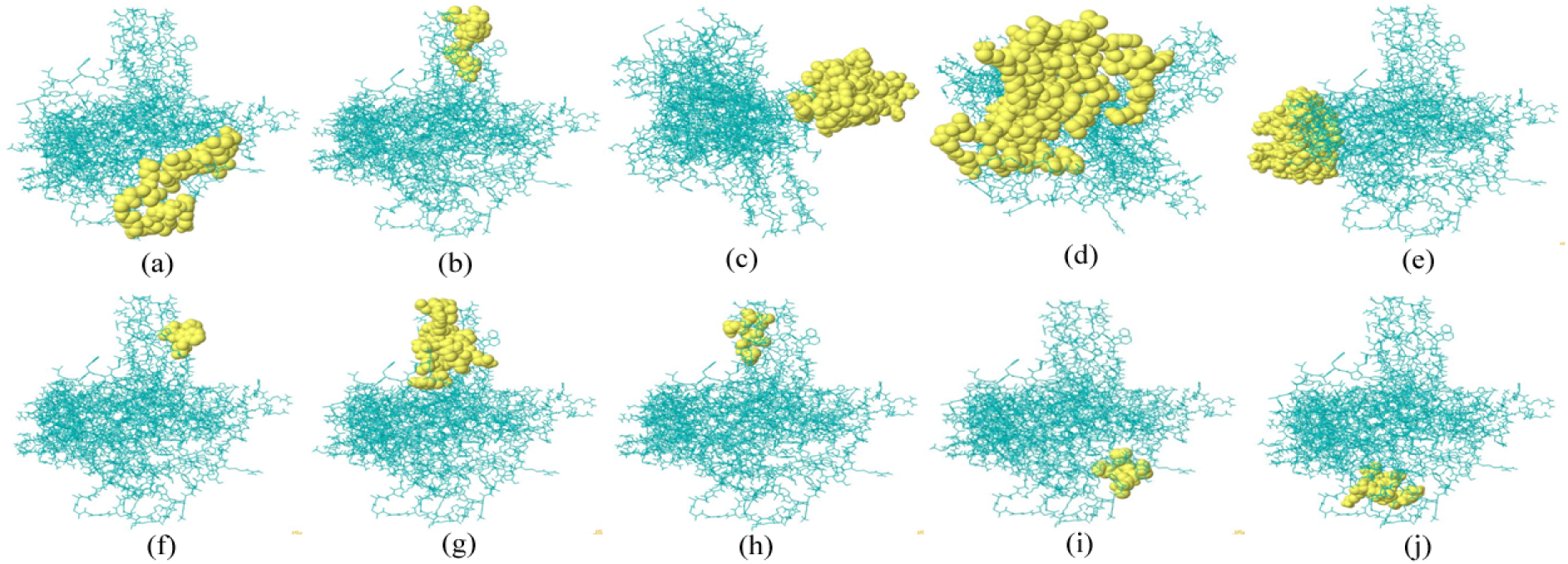
Ten conformational B-lymphocyte epitope mapping in final vaccine construct with length range 4-110 and score range 0.557-0.811 (cyan color represent the vaccine construct and yellow colored balls represent the conformational B-lymphocyte epitope).

### Molecular Docking of Final Vaccine Construct with TLR4 Immune Receptor

The vaccine-receptor docking was performed through ClusPro v2.0 for assessing the binding affinity of the vaccine construct with the immune receptor TLR4 and the ClusPro 2.0 server predicted a total of 26 complex forms (Tab. S4). Among them, the model having the lowest energy score properly occupied the receptor ^73^ and found that the model number 11 was selected as the best-docked complex (Fig. 10). The energy score of the best-docked complex was found to be −1292 which is the lowest among all other predicted complexes showing the highest binding interaction.

**Figure 10:**
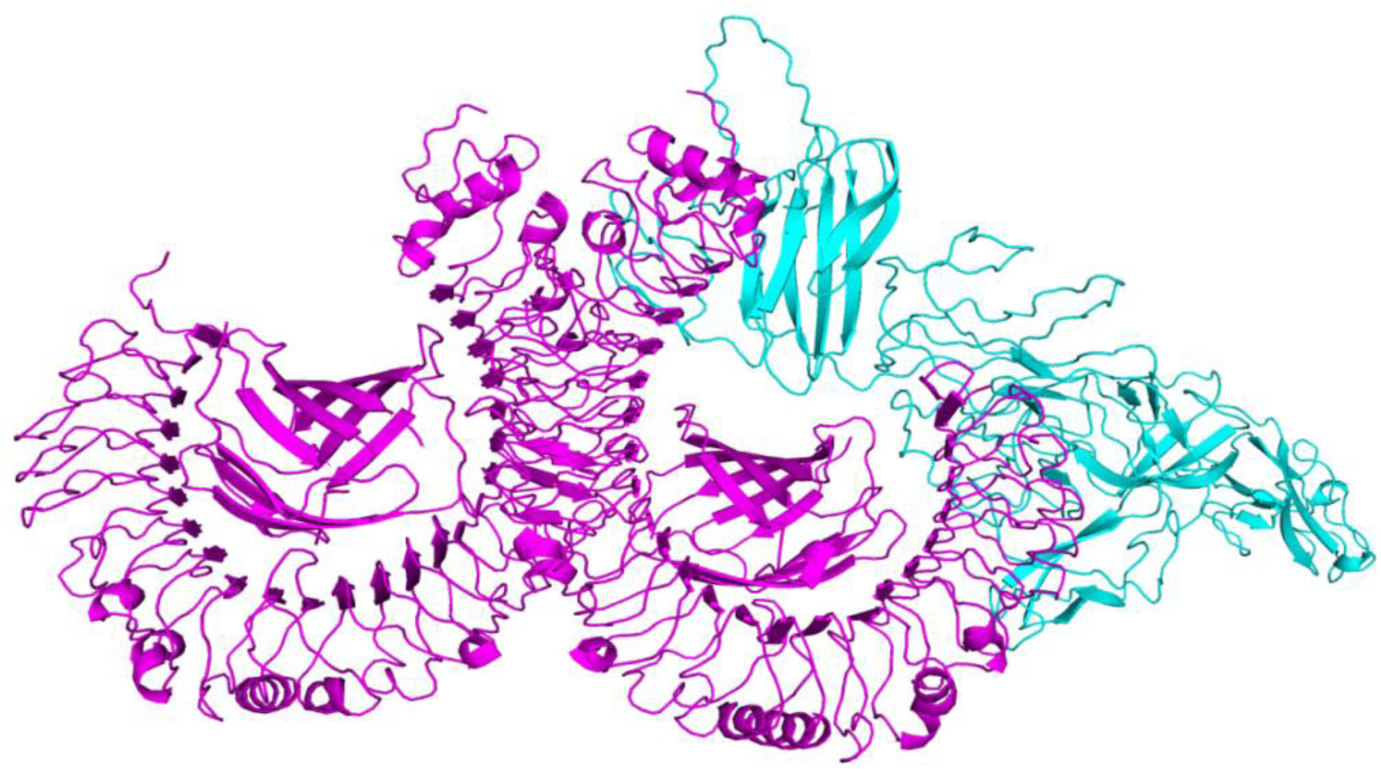
The docked complex of the vaccine construct with the TLR4 receptor (cyan color represent the vaccine construct and magenta color represent the TLR4 receptor).

### *In Silico* Cloning of the Final Vaccine Construct

To express the deigned vaccine protein into the *E. coli* expression system, *in silico* cloning was done. Therefore, it was necessary to adapt the codon respective to the vaccine constructs as per the codon usage of *E. coli* expression system. The adaptation of the codon usage of vaccine construct as per *E. coli* K12 was done using the JCat server. The JCat provided the optimized codon sequence of 1962 nucleotides, whose codon adaptive index (CAI) and percentage of the GC-content codon were found to be 0.98 and the percentage of the GC-content codon of 53.77%. The CAI value is very close to the best score 1.0 and the percentage of the GC-content codon lying in the allowed range 30-70%, therefore, both of them ensured the high expression rate of vaccine construct in *E. coli* K12 ^46^. Later on, the adapted codon sequence was inserted into the *E. coli* pET28a(+) vector and obtained a cloned vaccine of 7258 base pairs (Fig. 11).

**Figure 11:**
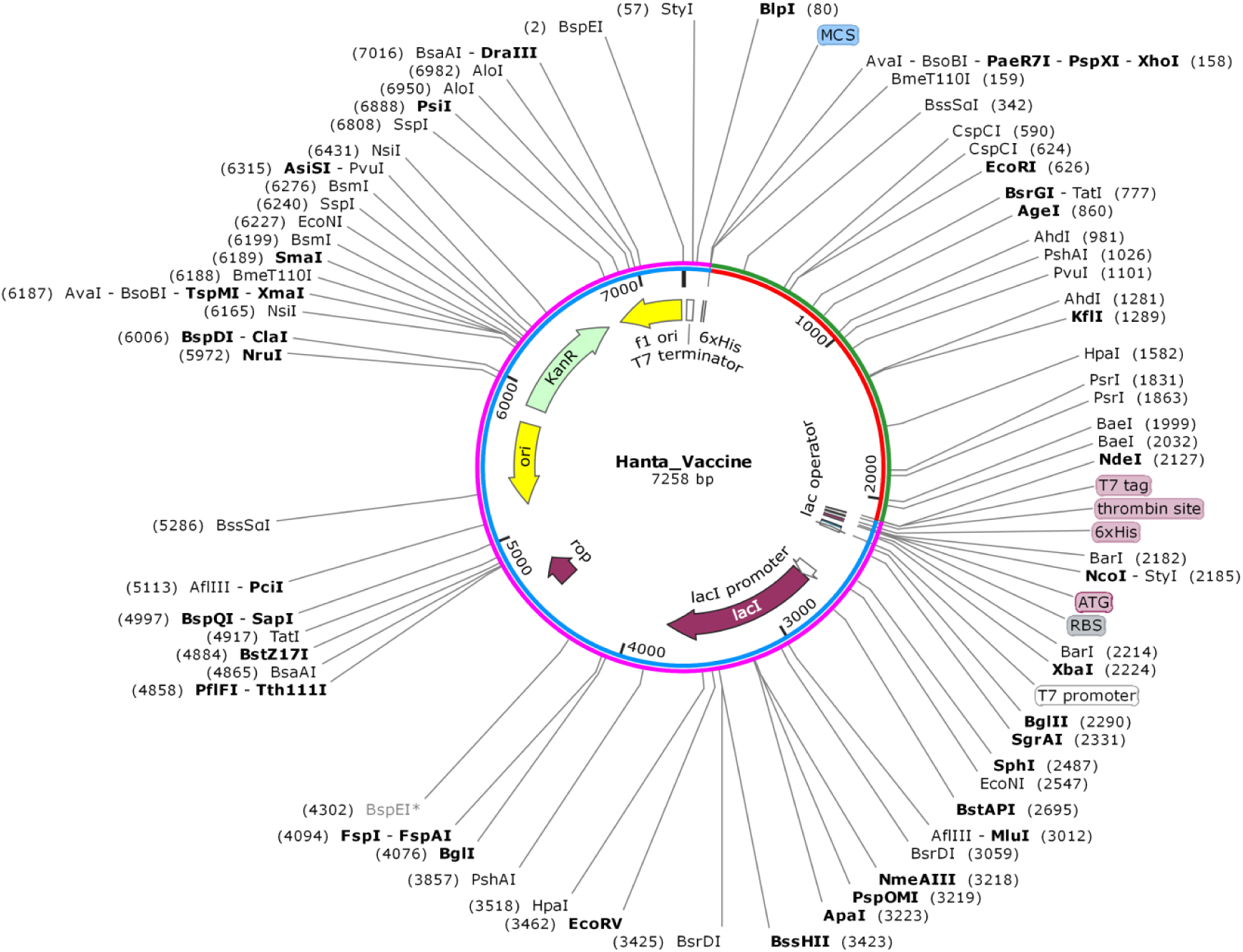
Cloned multi-epitope vaccine construct. *In silico* cloning of the adapted codon sequence of the final vaccine construct (red-green color) into the *E. coli* pET28a(+) vector (blue-magenta color).

### Disulfide Engineering of the Final Vaccine Construct

To stabilize the constructed vaccine structure, disulfide engineering was done through Disulfide by Design 2 (DbD2), and a total of 70 pairs of residues were predicted for the probable formation of disulfide bonds (Tab. S5). However, only 5 pairs of residues (Gly344-Ile507, Arg184-Pro265, Thr401-Gly436, Tyr413-Glu556, and Ala270-Pro375) were selected for the disulfide bond formation because their energy and Chi3 values lies within the allowed range i.e. the energy score should be less than 2.2 and the Chi3 value should be in between −87 to +97 ^74^. Thereafter, the stability of the vaccine structure for the mutation of every residue by cysteine in the selected 5 pairs was evaluated through DynaMut server and found that only three mutations of Gly344, Thr401, & Gly436 residues have positive vibrational entropy changes, which indicates the stability of the vaccine construct (Tab. S6). Among those three residues, Thr401 and Gly436 fulfill a pair for probable disulfide bond. Therefore, these two residues, Thr401 and Gly436 were taken into account for the mutation with cysteine (Fig. 12).

**Figure 12:**
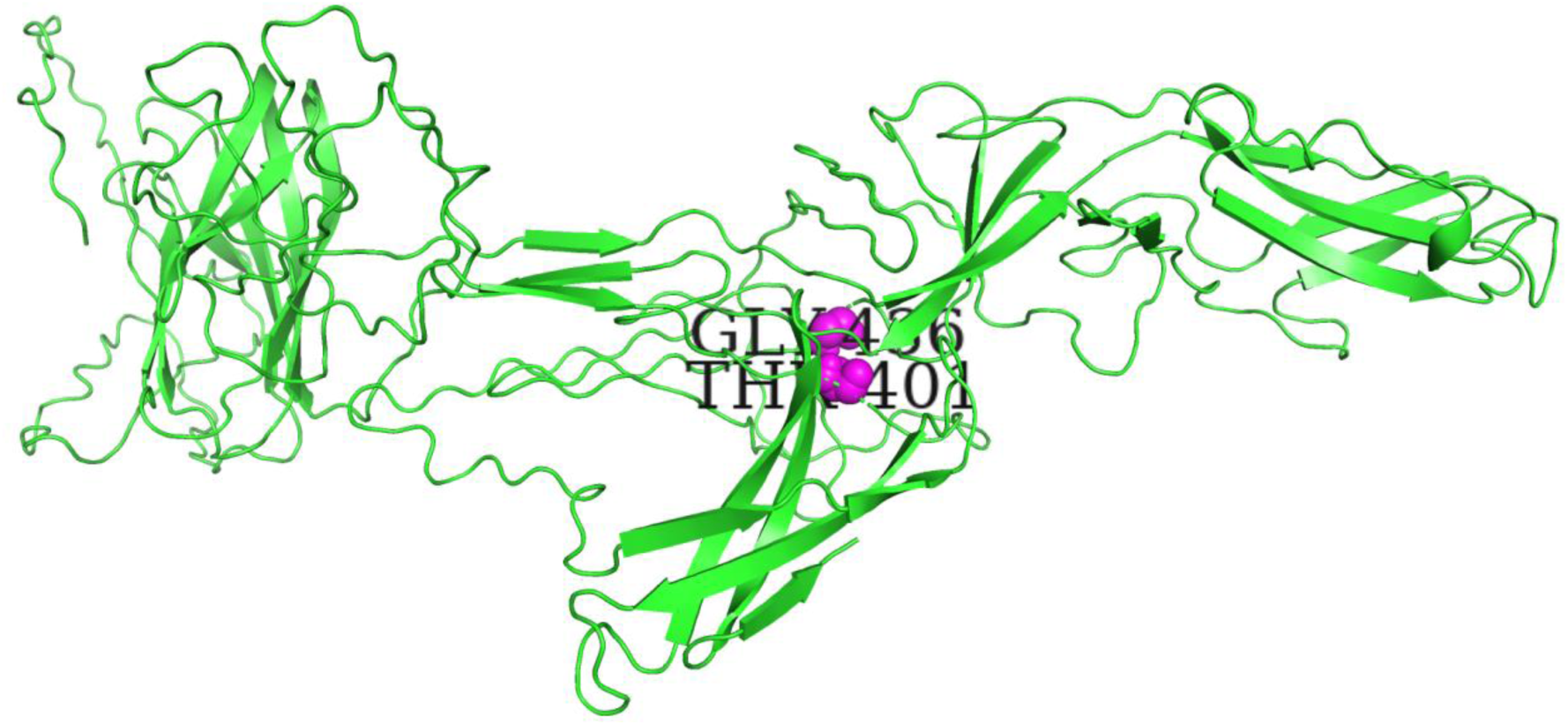
Disulfide engineering of the final vaccine construct where green color indicating the final vaccine construct and magenta colored ball indicating the pair of residues (Thr401-Gly436) for the probable formation of disulfide bond by mutating this pair with cysteines.

### Molecular Dynamics Simulation of the Best Vaccine-Receptor Docked Complex

The stability and large-scale mobility of the best vaccine-receptor docked complex were investigated by performing normal mode analysis (NMA) of its internal coordinates via iMODS. The vaccine protein and the receptor TLR4 were directed towards each other and the direction of each residue was represented by arrows (Fig. 13); where the length of the line indicates the degree of mobility. The vaccine-receptor complex deformability depends on the individual distortion of each residue, indicated by hinges in the high deformability region of the chain (Fig. 14(a)). The B-factor values inferred via NMA was equivalent to RMS (Fig. 14(b)). The complex eigenvalue was found to be 5.608279×10^−06^ (Fig. 14(c)); where the lower the eigenvalue, the easier the deformation ^75^. The eigenvalue is inversely related to the variance of the complex (Fig. 14(d)) ^76^. In the covariance matrix, the red, white, and blue colors indicate the correlated, uncorrelated, and anti-correlated pairs of residues (Fig. 14(e)) ^75,77^. An elastic network model was generated to represent the pair of atoms connected via springs (Fig. 14(f)). In the elastic graph, each dot represents one spring between the corresponding pair of atoms, where the darker grays indicate stiffer springs and vice versa ^75^.

**Figure 13:**
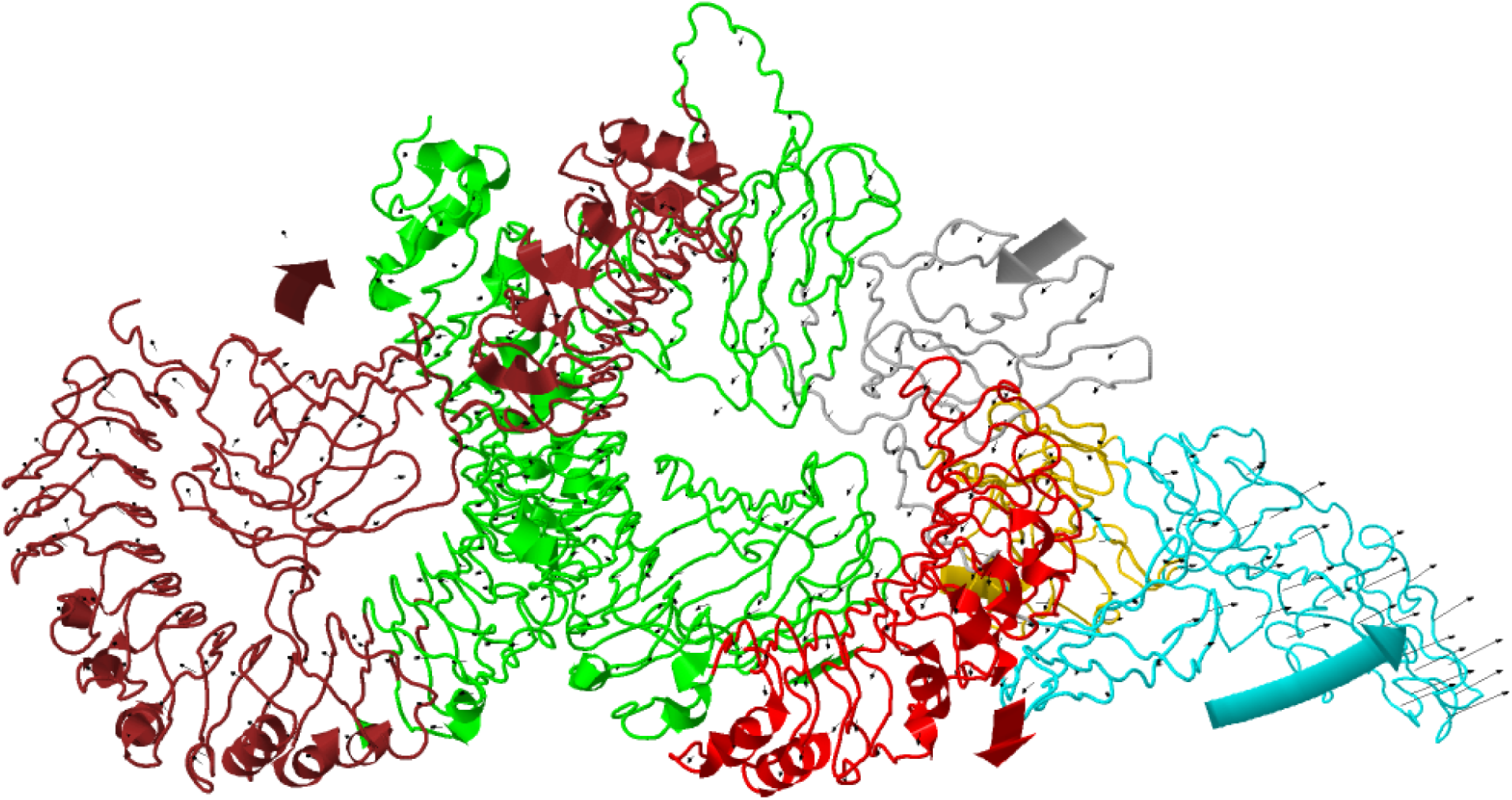
The vaccine-receptor docked complex after dynamics simulation. The large arrows indicate the direction of the vaccine protein and receptor TLR4 towards each other. The small arrows indicate the direction of the residues where the length of the line indicates the degree of mobility.

**Figure 14:**
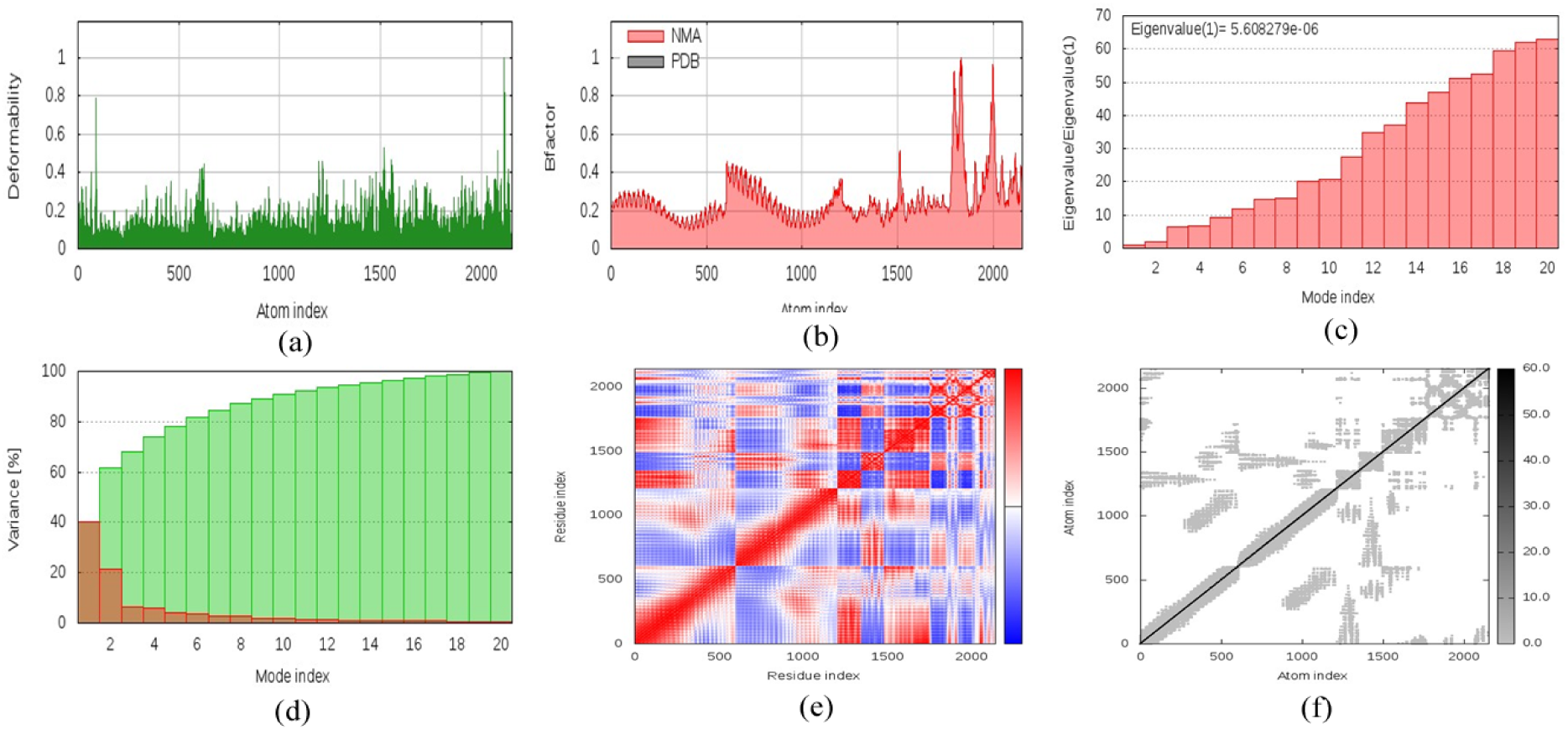
Molecular dynamics simulation of the best vaccine-receptor docked complex. (a, b) Deformability and B-factor at each atom, respectively. (c, d) Eigenvalue and variance of 20 modes of the docked complex, respectively. (e) The covariance matrix of the pairs of residues. (f) Elastic network graph of atoms.

### *In Silico* Immune Simulation of Final Vaccine Construct

The *in silico* immune response of the vaccine construct was generated by C-ImmSim immune simulator. The immune simulation showed results consistent with the typical immune response (Fig. 15). The primary response was identified by high levels of IgM. Following the secondary and tertiary responses were characterized by the high level of IgG1 + IgG2, IgM, and IgG + IgM antibodies with a corresponding decrease in antigen concentration (Fig. 15(a)).

**Figure 15:**
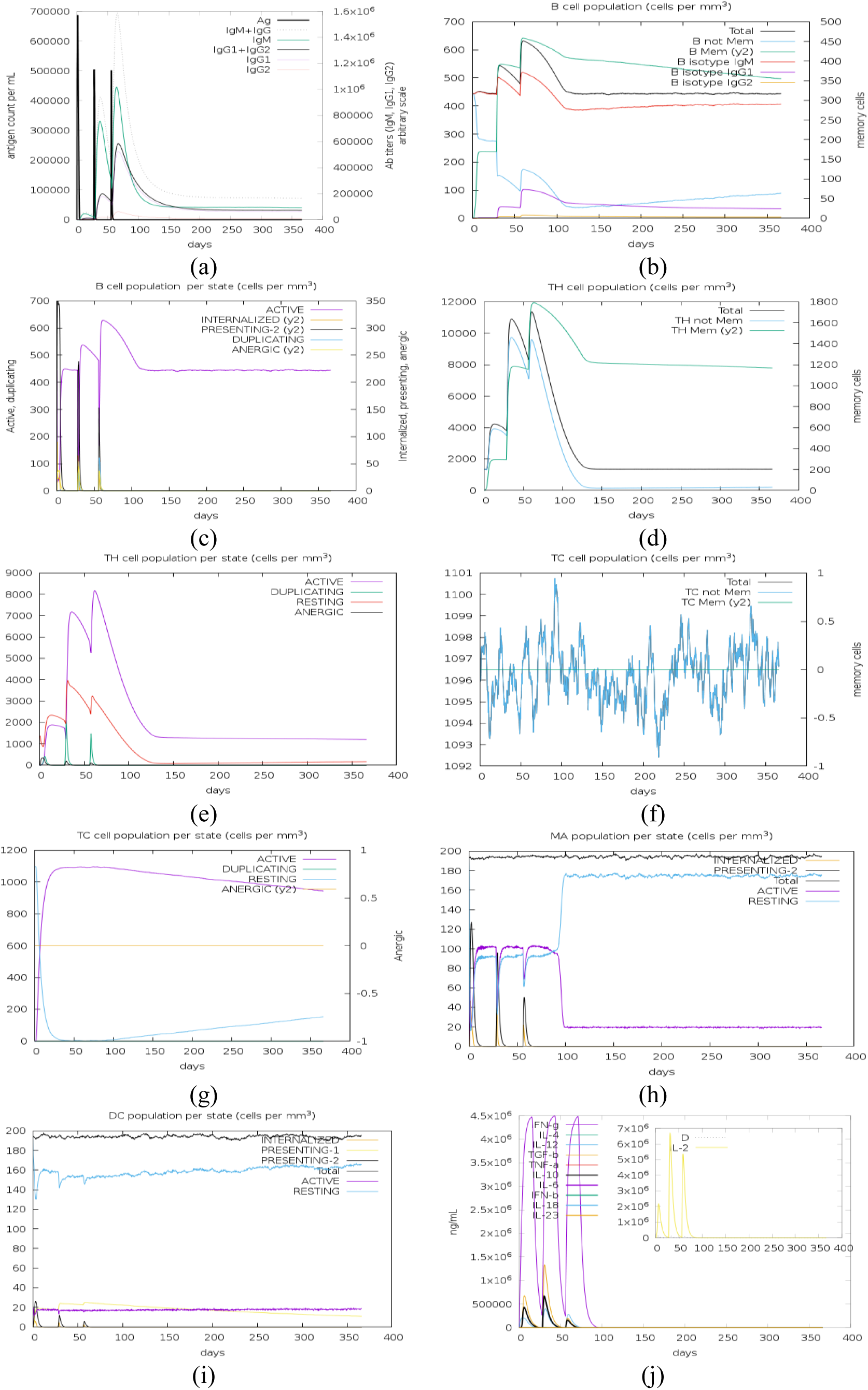
*In silico* simulation of immune response after injecting three injections by 4-weeks apart. (a) Antigen and immunoglobulins (antibodies are sub-divided per isotype). (b) B-lymphocytes count. (c) B-lymphocytes count per entity-state. (d) CD4 T-helper lymphocytes count. (e) CD4 T-helper lymphocytes count per entity-state. (f) CD8 T-cytotoxic lymphocytes count. (g) CD8 T-cytotoxic lymphocytes count per entity-state. (h) Macrophages count per entity-state. (i) Dendritic cell (DC) can present antigenic peptides on both MHC class-I and class-II molecules (the curves show the total number broken down to active, resting, internalized, and presenting the Ag). (j) Concentration of cytokines and interleukins. Simpson index (D) in the inset plot is danger signal.

Furthermore, several long-lasting B-cell isotypes were observed, indicating possible isotype switching potentials and memory formation (Fig. 15(b-c)). A similarly high response was highlighted in the T_H_ (helper) and T_C_ (cytotoxic) cell populations with respective memory development (Fig. 15(d-g)). During exposure, increased macrophage activity was demonstrated while dendritic cell activity was identified as consistent (Fig. 15(h-i)). The macrophages and dendritic cells are important when the expression of TLR4 in human is considered ^78^. High levels of IFN-γ and IL-2 were also evident and the lower the Simpson index (D), the greater the diversity (Fig. 15(j)) ^79^. The high level of IFN-γ and IL-2 production confirms the efficient Ig production, thereby, supporting a humoral response ^63^. In addition, there are many other characteristics have been produced such as PLB cell population, TR cell population per state, NK cell population, and EP population per state (Fig. S1); where all of them showed excellent immune response. This profile indicates the immune memory development and consequently the increased clearance of the antigen at subsequent exposures.

## Conclusion

The Zoonotic Hantavirus infection has emerged as a globally severe life-threatening problem characterized by an increasing number of infections and followed by deaths by 50%, worldwide. Despite this, there is no permanent cure or preventive treatment against Hantavirus is currently available. Therefore, it is urgent to explore an effective vaccine against Hantavirus to fight this severe problem. This study harnessed a complete series of immunoinformatics approaches to develop a novel multiepitope subunit vaccine containing T & B-cell epitopes for inducing cellular and humoral immunity, respectively. The T & B-cell epitopes were predicted from the selected highly antigenic proteins of every gene product. The predicted epitopes were filtered by antigenicity≥0.4 & immunogenicity>0 and followed by filtering by the non-toxin & non-allergic criterions. In addition, the helper T-cell epitopes were over-filtered by IL-10 & IFN-γ inducing criterions. Furthermore, the top conservancy associated epitope from every gene product was selected as potential vaccine candidate; among those the T-cell vaccine candidates along with their HLA alleles were subjected to analysis of the population coverage in both the epidemic and non-epidemic regions. Thereafter, the potential vaccine candidates were fused together with the help of suitable linkers for their adequate separate functions in the human body.

Subsequently, four vaccine models were generated by adding four different adjuvants to the nascent vaccine protein to increase the vaccine immunogenicity. The vaccine models were tested and compared for its antigenicity and allergenicity and followed by the structural analysis. Further, the physicochemical parameters of the final vaccine protein were evaluated and followed by the discontinuous B-cell epitope prediction. Molecular docking and dynamics simulation were also performed to investigate the binding affinity with immune receptor TLR-4 and stability of the vaccine-receptor complex, respectively and followed by conducting immune simulation to ensure the immune response and antigen clearance rate. At last, the disulfide engineering was performed to enhance the vaccine stability and followed by the *in silico* cloning to confirm the effective expression of the vaccine construct. However, this recommended vaccine requires experimental validation to ensure that it can induce immunity against Hantavirus.

## Methodology

### Retrieval of Whole Proteome, Grouping, and Assurance of Highest Antigenic Protein of Orthohantavirus

In order to design the subunit vaccine, the whole proteome of the *Orthohantavirus* was identified and found to be three gene products such as nucleoprotein, envelope glycoprotein (Gn, and Gcformerly known as G1, and G2, respectively), and RNA dependent RNA polymerase (RdRp) protein ^9^. The complete amino acid sequences of those proteins were retrieved from the NIAID Virus Pathogen Database and Analysis Resource (ViPR) ^80^ through the web site at www.viprbrc.org. The retrieved protein sequences were then grouped according to their strain, protein category, and epidemic nature. Thereafter the protein sequences of the targeted groups were subjected to the VaxiJen v2.0 server ^81^ available at http://www.ddg-pharmfac.net/vaxijen/VaxiJen/VaxiJen.html for predicting their antigenicity and ensuring the highest antigenic proteins for the final dataset preparation, where the threshold parameter was fixed as default.

### Prediction of Cytotoxic and Helper T-cell Epitopes with Corresponding Major Histocompatibility Complex Alleles and B-cell Epitopes

The cytotoxic T lymphocytes or CTLs or CD8+ T-cells are essential for immune defense against intracellular pathogens, including virus, bacteria, and tumour surveillance. Therefore, the highly antigenic proteins were submitted to the NetCTL v1.2 ^82^ server available at http://www.cbs.dtu.dk/services/NetCTL/ for predicting CD8+ T-cell epitopes. In the NetCTL server, we predicted the epitopes for 12 supertypes with a combined threshold score of 0.5. Furthermore, the IEDB MHC I Binding tool available at http://tools.iedb.org/mhci/ was used to predict the MHC-I binding alleles corresponding to each CD8+ T-cell epitopes ^83^; where the parameters consensus and human were set as the prediction method and MHC source species, respectively. In addition, the percentile rank≤2 was considered as alleles filtering criteria. On the other hand, the prediction of HTL or CD4+ T-cell epitopes is also a crucial stage of epitope-based vaccine design. The 15-mer CD4+ T-cell epitopes and their corresponding MHC-II binding alleles were predicted through the IEDB MHC II Binding tool (http://tools.iedb.org/mhcii/) using its consensus prediction method ^84^. A consensus percentile rank of ≤2 was set as a cutting point for allele selection because a lower percentile rank indicates a more binding affinity. B-cells are considered a core component of the adaptive immune system due to their ability to recognize and provide long-term protection against infectious pathogens by producing antibodies ^85^. Therefore, the prediction of B-cell epitopes is important for rational vaccine design. The highly antigenic protein sequences were subjected to the LBtope server (http://crdd.osdd.net/raghava/lbtope/) for predicting the B-cell epitopes ^86^. The LBtope predicts the B-cell epitopes based on the support vector machine (SVM) technique (the parameters are LBtope_Fixed, 20-mer peptide, and the percent of probability is 60%).

### Screening of CTL, HTL and LBL Epitopes for Potential Vaccine Candidates

The ability of CTL epitopes to trigger humoral and cell-mediated immune response called immunogenicity is the center of the vaccine efficacy ^87^. Consequently, the MHC I Immunogenicity tool of IEDB ^88^ available at http://tools.iedb.org/immunogenicity/ was used to analyze the immunogenicity of CTL epitopes. In the case of subunit vaccination, an epitope is considered to be effective if it has the ability (antigenicity) to induce protection from subsequent challenge by a disease-causing infective agent in an appropriate animal model following immunization ^81^. The epitope-based vaccine containing highly conserved epitopes can provide broader protection across multiple strains or even species ^89^. Therefore, the conservancy of the predicted epitopes was analyzed through the IEDB Conservation Across Antigens tool available at http://tools.iedb.org/conservancy/ ^89^. Another two important characteristics of peptides such as non-allergenicity and non-toxicity were checked using AllerTOP v2.0 ^67^ and ToxinPred server (http://crdd.osdd.net/raghava/toxinpred/) ^90^, respectively. The epitope-based vaccine should be designed to maximize population coverage in any geographical region especially in the region where the associated disease frequently occurs. In contrast, minimizing the number of T-cell epitopes included in the vaccine and the variability of population coverage obtained in different ethnic groups is important to be potential vaccine candidates ^91^. Hence, population coverage is a key factor for the development of the peptide-based vaccine ^92^. Finally, the IEDB Population Coverage tool ^91^ (http://tools.iedb.org/population/) was introduced to calculate the population coverage of the T-cell epitopes.

### Screening of Cytokine-inducing HTL Epitopes

Several cytokines such as IL-2, IL-6, IL-10, IFN-γ, and TNF-γ are associated with Hantavirus immune system ^93^. Herein, due to the tools unavailability, we only predicted the IL-10, and IFN-γ inducer HTL epitopes through the IL-10Pred server (https://webs.iiitd.edu.in/raghava/il10pred/) ^94^ and IFNepitope server (http://crdd.osdd.net/raghava/ifnepitope/) ^95^, respectively in which the SVM technique was chosen as a prediction method.

### Docking Simulation of the MHC HLA Alleles and the Peptide

The predominant binding affinities of the screened epitopes with their corresponding MHC HLA alleles of the known 3D structure were evaluated by the molecular docking simulation ^96^. Herein, we used PEPFOLD v3.5 server ^97^ to predict the tertiary structure of both the screened epitopes and the crystal structures of the lowest percentile ranked HLA proteins such as HLA-B*07:02 (6AT5), HLA-A*03:01 (2XPG), HLA-A*11:01 (5WJL), HLA-C*06:02 (5W6A), HLA-B*51:01 (1E27), HLA-B*39:01 (4O2F), HLA-A*24:02 (4F7M), HLA-B*40:02 (5IEK), HLA-B*18:01 (6MT3), & HLA-DRB1*01:01 (2FSE) ^98–107^. However, the allele structures were retrieved from Protein Data Bank (PDB). The Discovery Studio v16.0.0.400 was used for the separation of the protein and ligand and the AutoDock tools were used for the preparation of the protein and ligand files into PDBQT files by selecting the binding region and the number of torsion trees, respectively ^108^. Finally, these PDBQT files were analyzed by AutoDock Vina software for docking simulation ^109^.

### Cluster Analysis of the MHC Restricted HLA Alleles

In vaccine development, the resolved MHC super-families (clusters) play an important role for the identification of new targets with optimized affinity and selectivity of hits ^110^. The MHC HLA proteins with similar binding affinities were well clustered by the structure-based clustering technique ^110^. Herein, the MHCcluster v2.0 available at http://www.cbs.dtu.dk/services/MHCcluster/ was carried out to develop the heat-map of the HLAs functional relationship ^59^.

### Multi-Epitope Subunit Vaccine Design

An ideal effective multi-epitope vaccine should be composed of epitopes that can elicit CTL, HTL, and B cells and induce effective responses against the targeted virus ^39^. The screened CTL, HTL, and LBL vaccine candidates were conjugated in sequential manner to establish the final vaccine construct. In the vaccine construct, the epitopes were separated by proper linkers for their individual effective function ^111^. In this study, thelinkers GGGS, GPGPG, and KK were used for the separation of CTL, HTL, and LBL epitopes, respectively. Herein, the immunostimulatory adjuvants TLR4 agonist namely, RS-09, β3-defensin, β3-defensin with RR residues, and 50S ribosomal protein L7/L12 were used as adjuvants to construct four separate vaccine models. The problem of high polymorphic HLA alleles was solved by incorporating the PADRE sequence along with the adjuvants. The EAAAK linker was used to link the adjuvant with the CTL epitope.

### Immunological Assessment of the Vaccine Models

The antigenicity is a measure of the ability of an antigen to binds with the B & T-cell receptor that may lead to induce the immune response and memory cell formulation ^60^. Accordingly, the whole vaccine construct was submitted to the VaxiJen v2.0 server for predicting the antigenic nature of the vaccine protein. However, a vaccine with non-allergic behavior will be completely safe for human life ^70^. Therefore, the vaccine protein was analyzed for allergenicity using two modules, SVM (amino acid composition) and Hybrid approach under the AlgPred server (https://webs.iiitd.edu.in/raghava/algpred/submission.html) ^112^. The allergenicity of the vaccine protein was crosscheck through AllerTOP v2.0, and AllergenFP v1.0 ^113^.

### Secondary and Tertiary Structure Prediction of the Vaccine Models

The whole vaccine constructs were submitted to the Self-Optimized Prediction Method with Alignment (SOPMA) server (https://npsa-prabi.ibcp.fr/NPSA/npsa_sopma.html) for predicting the secondary structural features of the vaccine models ^114^. As a large portion of the immunoinformatics approach depends on the protein tertiary structure, the three-dimensional structure (3D) prediction is a crucial step for rational vaccine design. Therefore, the tertiary structures of the vaccine models were predicted using the RaptorX server available at http://raptorx.uchicago.edu/ ^69^. RaptorX is a multiple-template threading based protein modeling technique, which provides different factors of quality measure such as P-value for relative global quality, GDT & uGDT for absolute global quality, and modeling error at each residue ^69^.

### Refinement and Quality Assessment of the Tertiary Structure of the Vaccine Construct

The accuracy of the predicted tertiary structure is an important factor for protein docking, function and it depends on the degree of the alikeness of the target and available template structures ^46,115^. Although the template-based protein modeling is more accurate than the template-free protein modeling, but they may still have some problems in local and global structures and the errors such as irregular contacts, clashes, and unusual bond angles & lengths in the predicted 3D models ^115^. Since the inaccuracy in the predicted 3D models also limits the usage of the models for further studies, there was a necessity to improve the accuracy of the predicted model using refinement technique ^115^. The refinement of the predicted crude 3D model was done through the GalaxyRefine server (http://galaxy.seoklab.org/cgi-bin/submit.cgi?type=REFINE) ^68^. This server works based on a CASP10 tested refinement method which rebuilds the side chains and then repacking the overall structure by molecular dynamics simulation ^68^. To validate the accuracy of the refined 3D model, the Ramachandran plot was generated using PROCHECK under the PDBsum server available at www.ebi.ac.uk/pdbsum/ ^116,117^. The refined model also validated by calculating the quality Z-score using the ProSA-web server (https://prosa.services.came.sbg.ac.at/prosa.php) ^118^ and analyzing the non-bonded interactions using ERRAT server (http://servicesn.mbi.ucla.edu/ERRAT/) ^119^. The quality score outside the characteristic range of native proteins indicates possible errors in the predicted 3D structure.

### Physicochemical Properties Assessment of the Vaccine Construct

We further characterized the vaccine construct by assessing the various physicochemical properties including molecular weight (MW), theoretical pI, solubility, extinction coefficients (EC), estimated half-life, instability index (II), aliphatic index (AI) and grand average hydropathy (GRAVY) value associated with the vaccine construct. The physicochemical properties excluding solubility were computed by analyzing the vaccine protein sequence through the ProtParam tool under the bioinformatics resource portal Expert Protein Analysis System (ExPASy) (https://web.expasy.org/protparam/) ^72^. Later, the solubility of the vaccine protein upon over-expression in *E. coli* was predicted using the SOLpro tool in the SCRATCH suite ^120^.

### Conformational B–Cell epitopes Prediction of the Vaccine Construct

The patch of atoms on the protein surface is termed as the discontinuous or conformational epitope. It is predicted to design a molecule that can mimic the structure and immunogenic properties of an epitope and replace it either in the process of antibody production-in this case an epitope mimic can be considered as a prophylactic or therapeutic vaccine-or antibody detection in medical diagnostics or experimental research ^121^. Therefore, the CBLs of the vaccine construct were predicted by subjecting the 3D structure of the vaccine construct to the IEDB ElliPro tool available at http://tools.iedb.org/ellipro/ ^121^.

### Disulfide Engineering of Vaccine Construct

Disulfide bonds can provide increased stability of the protein by reducing unfolded conformational entropy and raising the free energy of the denatured state along with the examination of protein interactions and dynamics ^70^. The Disulfide by Design 2 (DbD2) v2.12 server available at http://cptweb.cpt.wayne.edu/DbD2/ was carried out to perform the disulfide engineering of the predicted 3D model of the vaccine construct ^74^. The DbD2 can predict the probable pairs of residues that have the characteristics to forma disulfide bond by mutating the individual residues with cysteine. Afterward, the DynaMut server available at http://biosig.unimelb.edu.au/dynamut/ was used to crosscheck the stability of the vaccine construct for the mutation by cysteine of each residue predicted by DbD2 ^122^. The DynaMut server predicts the stability based on the vibrational entropy changes, where the positive value indicates the stability of the vaccine construct.

### Molecular Docking of Vaccine Construct with Immunological Receptor Protein

Molecular docking helps us to characterize the small molecules (ligand) in the binding site of target proteins (receptor) by modeling the ligand-receptor complex structure at the atomic level and predicting the tentative binding parameters of ligand-receptor complex ^123^. The host endothelial cells become dysfunctional and pathogenic by the innate immune response induced by Hantavirus ^124^. As toll-like receptor 4 (TLR4) expression is upregulated and mediates the secretion of several cytokines in Hantaan virus infected endothelial cells, it was used as an immunological receptor protein to induce anti-hantavirus immunity ^124^. The TLR4 (PDB ID: 4G8A) was retrieved from RCSB-Protein Data Bank. The vaccine construct was treated as ligand protein. The ClusPro 2.0 server available at https://cluspro.bu.edu/login.php helps to finish the docking of vaccine construct with the TLR4 receptor protein ^125^. The docked complex with the lowest energy score was selected as the best complex.

### Molecular Dynamics Simulation of Vaccine-Receptor Complex

The molecular dynamic simulation is the most popular computational simulation approach for studying the physical basis of the structure and function of the biological macromolecules ^126^. The dynamic simulations with biologically relevant sizes and time scales are difficult for understanding macromolecular functioning ^127^. The comparison of the essential dynamics of proteins to their normal modes is an alternative powerful tool than the costly atomistic simulation ^127–129^. The stability of the vaccine-receptor complex was determined through the iMOD server (iMODS) available at http://imods.chaconlab.org/, which describes the collective functional motions in terms of deformability, eigenvalues, and covariance map by analyzing normal modes in internal coordinates ^75^. The capacity of a given molecule to deform at each of its residues is measured by the deformability. The eigenvalue associated with each normal mode indicates the motion stiffness and it is directly related to the energy required to deform the structure. The covariance map represents the coupling between pairs of residues.

### Immune Simulation of Vaccine Construct

The immune simulation is a key step for understanding the immune system by determining the immunogenicity and immune response profile of the vaccine protein ^79^. The agent-based immune simulator, C-ImmSim available at http://kraken.iac.rm.cnr.it/C-IMMSIM/ was utilized to conduct the immune simulation of the vaccine protein. The C-ImmSim simulator uses the position-specific scoring matrix (PSSM) and machine learning techniques for the prediction of immune epitopes and immune interactions, respectively ^79^. The minimum recommended time interval between doses is four weeks for most of the currently used vaccines and in accordance with the TOVA approach; three injections are given at intervals of four weeks for the target product profile of a prophylactic onchocerciasis vaccine ^63,130^. Therefore, three injections of the designed peptide were considered at an interval of four weeks with conserved host HLA alleles and time step set at 1, 84, and 168 (each time step is equivalent to 8 hours of real-life and time step 1 is injection at time=0). The simulation volume and simulation steps were set at 110 and 1100, respectively. The remaining simulation parameters were set at default (random seed=12345, what to inject=vaccine (no LPS), adjuvant=100, and num Ag to inject=1000). The Simpson index, D (a measure of diversity) was interpreted from the plot.

### Codon Adaptation and *In Silico* Cloning of Vaccine Construct

Codon adaptation is a process to increase the expression rate of the foreign genes in the host when the codon usages in both organisms differ from each other. Therefore, the Java Codon Adaptation Tool (JCat) available at http://www.jcat.de/ was used to express the vaccine protein in a most sequenced prokaryotic organism, *E. coli* K12 ^73,131^. The expression rate parameters, the codon adaptation index (CAI) and the percentage of GC-content codon should be in range 0.8-1.0 and 30-70%, respectively, where CAI of 1.0 is considered as the best score ^46^. In JCat, additional three options were checked to avoid rho-independent transcription terminators, prokaryotic ribosome binding sites, and cleavage sites of restriction enzymes. Furthermore, in the optimized nucleotide sequence, the absence of the restriction enzyme cutting sites (XhoI and NdeI) was confirmed. Afterward, the XhoI and NdeI restriction sites were added to the N and C-terminal of the optimized sequence, respectively. Finally, the restriction cloning module of the SnapGene tool was used to clone the optimized adapted codon sequence of the vaccine construct into the *E. coli* pET28a(+) vector.

## Competing interests

The author(s) have no competing interests.

## Author contributions

FA and UKA planned & designed the protocol; FA and UKA prepared the data; FA and ZN performed the computational analysis; FA prepared all the results; FA and UKA wrote the manuscript; FA, SBS and MSAK contributed to screen the vaccine candidates; MMH contributed to the critical revision of the manuscript; UKA supervised the whole study.

## References

1. Kruger, D. H., Figueiredo, L. T. M., Song, J.-W. & Klempa, B. Hantaviruses—Globally emerging pathogens. J. Clin. Virol. 64, 128–136 (2015).

2. Heinemann, P. et al. Human Infections by Non–Rodent-Associated Hantaviruses in Africa. J. Infect. Dis. 214, 1507–1511 (2016).

3. Jiang, H. et al. Hantavirus infection: a global zoonotic challenge. Virol. Sin. 32, 32–43 (2017).

4. Zhang, Y.-Z. Discovery of hantaviruses in bats and insectivores and the evolution of the genus Hantavirus. Virus Res. 187, 15–21 (2014).

5. Guo, W.-P. et al. Phylogeny and Origins of Hantaviruses Harbored by Bats, Insectivores, and Rodents. PLOS Pathog. 9, e1003159 (2013).

6. Reynes, J.-M. et al. Anjozorobe hantavirus, a new genetic variant of Thailand virus detected in rodents from Madagascar. Vector Borne Zoonotic Dis. 14, 212–219 (2014).

7. Zhang, W.-Y. et al. Spatiotemporal Transmission Dynamics of Hemorrhagic Fever with Renal Syndrome in China, 2005–2012. PLoS Negl. Trop. Dis. 8, e3344 (2014).

8. Schmaljohn, C. & Hjelle, B. Hantaviruses: a global disease problem. Emerg. Infect. Dis. 3, 95–104 (1997).

9. Jonsson, C. B., Figueiredo, L. T. M. & Vapalahti, O. A global perspective on hantavirus ecology, epidemiology, and disease. Clin. Microbiol. Rev. 23, 412–441 (2010).

10. Jiang, W. et al. Development of a SYBR Green I based one-step real-time PCR assay for the detection of Hantaan virus. J. Virol. Methods 196, 145–151 (2014).

11. Zou, L.-X., Chen, M.-J. & Sun, L. Haemorrhagic fever with renal syndrome: literature review and distribution analysis in China. Int. J. Infect. Dis. 43, 95–100 (2016).

12. Jiang, H., Du, H., Wang, L. M., Wang, P. Z. & Bai, X. F. Hemorrhagic Fever with Renal Syndrome: Pathogenesis and Clinical Picture. Front. Cell. Infect. Microbiol. 6, 1 (2016).

13. Zhang, S. et al. Epidemic characteristics of hemorrhagic fever with renal syndrome in China, 2006–2012. BMC Infect. Dis. 14, 384 (2014).

14. Papa, A. et al. Meeting report: Tenth International Conference on Hantaviruses. Antiviral Res. 133, 234–241 (2016).

15. Onishchenko, G. & Ezhlova, E. Epidemiologic surveillance and prophylaxis of hemorrhagic fever with renal syndrome in Russian Federation. Zh. Mikrobiol. Epidemiol. Immunobiol. 23–32 (2013).

16. Krüger, D. H., Ulrich, R. G. & Hofmann, J. Hantaviruses as zoonotic pathogens in Germany. Dtsch. Arztebl. Int. 110, 461–467 (2013).

17. Demeester, R. et al. Hantavirus nephropathy as a pseudo-import pathology from Ecuador. Eur. J. Clin. Microbiol. Infect. Dis. 29, 59 (2009).

18. Knust, B. & Rollin, P. E. Twenty-year summary of surveillance for human hantavirus infections, United States. Emerg. Infect. Dis. 19, 1934–1937 (2013).

19. Goeijenbier, M. et al. Emerging Viruses in the Republic of Suriname: Retrospective and Prospective Study into Chikungunya Circulation and Suspicion of Human Hantavirus Infections, 2008–2012 and 2014. Vector-Borne Zoonotic Dis. 15, 611–618 (2015).

20. Figueiredo, L. T. M., Souza, W. M. de, Ferrés, M. & Enria, D. A. Hantaviruses and cardiopulmonary syndrome in South America. Virus Res. 187, 43–54 (2014).

21. Armien, B. et al. Hantavirus fever without pulmonary syndrome in Panama. Am. J. Trop. Med. Hyg. 89, 489–494 (2013).

22. Lee, H. W., Lee, P. W. & Johnson, K. M. Isolation of the Etiologic Agent of Korean Hemorrhagic Fever. J. Infect. Dis. 137, 298–308 (1978).

23. Latus, J. et al. Acute kidney injury and tools for risk-stratification in 456 patients with hantavirus-induced nephropathia epidemica. Nephrol. Dial. Transplant. 30, 245–251 (2014).

24. Brummer-Korvenkontio, M. et al. Nephropathia Epidemica: Detection of Antigen in Bank Voles and Serologic Diagnosis of Human Infection. J. Infect. Dis. 141, 131–134 (1980).

25. Vollmar, P. et al. Hantavirus cardiopulmonary syndrome due to Puumala virus in Germany. J. Clin. Virol. 84, 42–47 (2016).

26. Noh, J. Y. et al. Clinical and molecular epidemiological features of hemorrhagic fever with renal syndrome in Korea over a 10-year period. J. Clin. Virol. 58, 11–17 (2013).

27. Yao, L.-S. et al. Complete genome sequence of Seoul virus isolated from Rattus norvegicus in the Democratic People’s Republic of Korea. J. Virol. 86, 13853 (2012).

28. Papa, A. Dobrava-Belgrade virus: Phylogeny, epidemiology, disease. Antiviral Res. 95, 104–117 (2012).

29. Nikolic, V. et al. Evidence of recombination in Tula virus strains from Serbia. Infect. Genet. Evol. 21, 472–478 (2014).

30. Pattamadilok, S. et al. GEOGRAPHICAL DISTRIBUTION OF HANTAVIRUSES IN THAILAND AND POTENTIAL HUMAN HEALTH SIGNIFICANCE OF THAILAND VIRUS. Am. J. Trop. Med. Hyg. 75, 994–1002 (2006).

31. Gamage, C. D. et al. Serological evidence of Thailand virus-related hantavirus infection among suspected leptospirosis patients in Kandy, Sri Lanka. Jpn. J. Infect. Dis. 64, 72–75 (2011).

32. Raharinosy, V. et al. Geographical distribution and relative risk of Anjozorobe virus (Thailand orthohantavirus) infection in black rats (Rattus rattus) in Madagascar. Virol. J. 15, 83 (2018).

33. Klempa, B. et al. Serological Evidence of Human Hantavirus Infections in Guinea, West Africa. J. Infect. Dis. 201, 1031–1034 (2010).

34. Ksiazek, T. G. et al. Identification of a New North American Hantavirus that Causes Acute Pulmonary Insufficiency. Am. J. Trop. Med. Hyg. 52, 117d–123 (1995).

35. Krüger, D. H., Schönrich, G. & Klempa, B. Human pathogenic hantaviruses and prevention of infection. Hum. Vaccin. 7, 685–693 (2011).

36. Avšič-Županc, T., Saksida, A. & Korva, M. Hantavirus infections. Clin. Microbiol. Infect. 21, e6–e16 (2019).

37. Hinrichsen, S. L. & Godoi, J. T. Hantavirus disease. Rev. Assoc. Med. Bras. 42, 237–241 (1996).

38. Calisher, C. H. et al. EPIZOOTIOLOGY OF SIN NOMBRE AND EL MORO CANYON HANTAVIRUSES, SOUTHEASTERN COLORADO, 1995–2000. J. Wildl. Dis. 41, 1–11 (2005).

39. Zhang, L. Multi-epitope vaccines: a promising strategy against tumors and viral infections. Cell. Mol. Immunol. 15, 182–184 (2018).

40. Buonaguro, L. & Consortium, H. Developments in cancer vaccines for hepatocellular carcinoma. Cancer Immunol. Immunother. 65, 93–99 (2016).

41. Brennick, C. A., George, M. M., Corwin, W. L., Srivastava, P. K. & Ebrahimi-Nik, H. Neoepitopes as cancer immunotherapy targets: key challenges and opportunities. Immunotherapy 9, 361–371 (2017).

42. Kuo, T., Wang, C., Badakhshan, T., Chilukuri, S. & BenMohamed, L. The challenges and opportunities for the development of a T-cell epitope-based herpes simplex vaccine. Vaccine 32, 6733–6745 (2014).

43. He, R. et al. Efficient control of chronic LCMV infection by a CD4 T cell epitope-based heterologous prime-boost vaccination in a murine model. Cell. Mol. Immunol. 15, 815–826 (2018).

44. Lu, I.-N., Farinelle, S., Sausy, A. & Muller, C. P. Identification of a CD4 T-cell epitope in the hemagglutinin stalk domain of pandemic H1N1 influenza virus and its antigen-driven TCR usage signature in BALB/c mice. Cell. Mol. Immunol. 14, 511–520 (2017).

45. Abdulla, F., Adhikari, U. K. & Uddin, M. K. Exploring T & B-cell epitopes and designing multi-epitope subunit vaccine targeting integration step of HIV-1 lifecycle using immunoinformatics approach. Microb. Pathog. 137, 103791 (2019).

46. Ali, M. et al. Exploring dengue genome to construct a multi-epitope based subunit vaccine by utilizing immunoinformatics approach to battle against dengue infection. Sci. Rep. 7, 9232 (2017).

47. Depla, E. et al. Rational design of a multiepitope vaccine encoding T-lymphocyte epitopes for treatment of chronic hepatitis B virus infections. J. Virol. 82, 435–450 (2008).

48. Nosrati, M., Mohabatkar, H. & Behbahani, M. A Novel Multi-Epitope Vaccine For Cross Protection Against Hepatitis C Virus (HCV): An Immunoinformatics Approach TT -. Res-Mol-Med 5, 17–26 (2017).

49. Ikram, A. et al. Exploring NS3/4A, NS5A and NS5B proteins to design conserved subunit multi-epitope vaccine against HCV utilizing immunoinformatics approaches. Sci. Rep. 8, 16107 (2018).

50. Bazhan, S. I. et al. In silico Designed Ebola Virus T-Cell Multi-Epitope DNA Vaccine Constructions Are Immunogenic in Mice. Vaccines 7, 34 (2019).

51. Narula, A., Pandey, R. K., Khatoon, N., Mishra, A. & Prajapati, V. K. Excavating chikungunya genome to design B and T cell multi-epitope subunit vaccine using comprehensive immunoinformatics approach to control chikungunya infection. Infect. Genet. Evol. 61, 4–15 (2018).

52. Hasan, M. et al. Reverse vaccinology approach to design a novel multi-epitope subunit vaccine against avian influenza A (H7N9) virus. Microb. Pathog. 130, 19–37 (2019).

53. Kumar Pandey, R., Ojha, R., Mishra, A. & Kumar Prajapati, V. Designing B- and T-cell multi-epitope based subunit vaccine using immunoinformatics approach to control Zika virus infection. J. Cell. Biochem. 119, 7631–7642 (2018).

54. Ying, J., Dong, X. & Chen, Y. Preliminary evaluation of a candidate multi-epitope-vaccine against the classical swine fever virus. Tsinghua Sci. Technol. 13, 433–438 (2008).

55. Ojha, R., Pareek, A., Pandey, R. K., Prusty, D. & Prajapati, V. K. Strategic Development of a Next-Generation Multi-Epitope Vaccine To Prevent Nipah Virus Zoonotic Infection. ACS omega 4, 13069–13079 (2019).

56. Azim, K. F. et al. Immunoinformatics approaches for designing a novel multi epitope peptide vaccine against human norovirus (Norwalk virus). Infect. Genet. Evol. 74, 103936 (2019).

57. Adhikari, U. K. & Rahman, M. M. Overlapping CD8+ and CD4+ T-cell epitopes identification for the progression of epitope-based peptide vaccine from nucleocapsid and glycoprotein of emerging Rift Valley fever virus using immunoinformatics approach. Infect. Genet. Evol. 56, 75–91 (2017).

58. Adhikari, U. K., Tayebi, M. & Rahman, M. M. Immunoinformatics Approach for Epitope-Based Peptide Vaccine Design and Active Site Prediction against Polyprotein of Emerging Oropouche Virus. J. Immunol. Res. 2018, 6718083 (2018).

59. Thomsen, M., Lundegaard, C., Buus, S., Lund, O. & Nielsen, M. MHCcluster, a method for functional clustering of MHC molecules. Immunogenetics 65, 655–665 (2013).

60. Pandey, R. K., Bhatt, T. K. & Prajapati, V. K. Novel Immunoinformatics Approaches to Design Multi-epitope Subunit Vaccine for Malaria by Investigating Anopheles Salivary Protein. Sci. Rep. 8, 1125 (2018).

61. Rahmani, A. et al. Development of a conserved chimeric vaccine based on helper T-cell and CTL epitopes for induction of strong immune response against Schistosoma mansoni using immunoinformatics approaches. Int. J. Biol. Macromol. 141, 125–136 (2019).

62. Kalita, P., Lyngdoh, D. L., Padhi, A. K., Shukla, H. & Tripathi, T. Development of multi-epitope driven subunit vaccine against Fasciola gigantica using immunoinformatics approach. Int. J. Biol. Macromol. 138, 224–233 (2019).

63. Shey, R. A. et al. In-silico design of a multi-epitope vaccine candidate against onchocerciasis and related filarial diseases. Sci. Rep. 9, 4409 (2019).

64. Shanmugam, A. et al. Synthetic Toll Like Receptor-4 (TLR-4) Agonist Peptides as a Novel Class of Adjuvants. PLoS One 7, e30839 (2012).

65. Mohan, T., Sharma, C., Bhat, A. A. & Rao, D. N. Modulation of HIV peptide antigen specific cellular immune response by synthetic α- and β-defensin peptides. Vaccine 31, 1707–1716 (2013).

66. Lee, S. J. et al. A Potential Protein Adjuvant Derived from Mycobacterium tuberculosis Rv0652 Enhances Dendritic Cells-Based Tumor Immunotherapy. PLoS One 9, e104351 (2014).

67. Dimitrov, I., Flower, D. R. & Doytchinova, I. AllerTOP--a server for in silico prediction of allergens. BMC Bioinformatics 14 Suppl 6, S4–S4 (2013).

68. Heo, L., Park, H. & Seok, C. GalaxyRefine: Protein structure refinement driven by side-chain repacking. Nucleic Acids Res. 41, W384–W388 (2013).

69. Källberg, M. et al. Template-based protein structure modeling using the RaptorX web server. Nat. Protoc. 7, 1511–1522 (2012).

70. Khatoon, N., Pandey, R. K. & Prajapati, V. K. Exploring Leishmania secretory proteins to design B and T cell multi-epitope subunit vaccine using immunoinformatics approach. Sci. Rep. 7, 8285 (2017).

71. Naz, A. et al. Identification of putative vaccine candidates against Helicobacter pylori exploiting exoproteome and secretome: A reverse vaccinology based approach. Infect. Genet. Evol. 32, 280–291 (2015).

72. Wilkins, M. R. et al. Protein Identification and Analysis Tools in the ExPASy Server BT - 2-D Proteome Analysis Protocols. in (ed. Link, A. J.) 531–552 (Humana Press, 1999). doi:10.1385/1-59259-584-7:531

73. Pandey, R. K., Ojha, R., Aathmanathan, V. S., Krishnan, M. & Prajapati, V. K. Immunoinformatics approaches to design a novel multi-epitope subunit vaccine against HIV infection. Vaccine 36, 2262–2272 (2018).

74. Craig, D. B. & Dombkowski, A. A. Disulfide by Design 2.0: a web-based tool for disulfide engineering in proteins. BMC Bioinformatics 14, 346 (2013).

75. López-Blanco, J. R., Aliaga, J. I., Quintana-Ortí, E. S. & Chacón, P. iMODS: internal coordinates normal mode analysis server. Nucleic Acids Res. 42, W271–W276 (2014).

76. Kovacs, J. A., Chacón, P. & Abagyan, R. Predictions of protein flexibility: First-order measures. Proteins Struct. Funct. Bioinforma. 56, 661–668 (2004).

77. Ichiye, T. & Karplus, M. Collective motions in proteins: A covariance analysis of atomic fluctuations in molecular dynamics and normal mode simulations. Proteins Struct. Funct. Bioinforma. 11, 205–217 (1991).

78. Duthie, M. S., Windish, H. P., Fox, C. B. & Reed, S. G. Use of defined TLR ligands as adjuvants within human vaccines. Immunol. Rev. 239, 178–196 (2011).

79. Rapin, N., Lund, O., Bernaschi, M. & Castiglione, F. Computational Immunology Meets Bioinformatics: The Use of Prediction Tools for Molecular Binding in the Simulation of the Immune System. PLoS One 5, e9862 (2010).

80. Pickett, B. E. et al. ViPR: an open bioinformatics database and analysis resource for virology research. Nucleic Acids Res. 40, D593–D598 (2012).

81. Doytchinova, I. A. & Flower, D. R. VaxiJen: a server for prediction of protective antigens, tumour antigens and subunit vaccines. BMC Bioinformatics 8, 4 (2007).

82. Larsen, M. V et al. Large-scale validation of methods for cytotoxic T-lymphocyte epitope prediction. BMC Bioinformatics 8, 424 (2007).

83. Moutaftsi, M. et al. A consensus epitope prediction approach identifies the breadth of murine TCD8+-cell responses to vaccinia virus. Nat. Biotechnol. 24, 817–819 (2006).

84. Wang, P. et al. Peptide binding predictions for HLA DR, DP and DQ molecules. BMC Bioinformatics 11, 568 (2010).

85. Jespersen, M. C., Peters, B., Nielsen, M. & Marcatili, P. BepiPred-2.0: improving sequence-based B-cell epitope prediction using conformational epitopes. Nucleic Acids Res. 45, W24–W29 (2017).

86. Singh, H., Ansari, H. R. & Raghava, G. P. S. Improved Method for Linear B-Cell Epitope Prediction Using Antigen’s Primary Sequence. PLoS One 8, e62216 (2013).

87. Leroux-Roels, G., Bonanni, P., Tantawichien, T. & Zepp, F. Vaccine development. Perspect. Vaccinol. 1, 115–150 (2011).

88. Calis, J. J. A. et al. Properties of MHC class I presented peptides that enhance immunogenicity. PLoS Comput. Biol. 9, e1003266–e1003266 (2013).

89. Bui, H.-H., Sidney, J., Li, W., Fusseder, N. & Sette, A. Development of an epitope conservancy analysis tool to facilitate the design of epitope-based diagnostics and vaccines. BMC Bioinformatics 8, 361 (2007).

90. Gupta, S. et al. In Silico Approach for Predicting Toxicity of Peptides and Proteins. PLoS One 8, e73957 (2013).

91. Bui, H.-H. et al. Predicting population coverage of T-cell epitope-based diagnostics and vaccines. BMC Bioinformatics 7, 153 (2006).

92. Khan, M. A., Hossain, M. U., Rakib-Uz-Zaman, S. M. & Morshed, M. N. Epitope-based peptide vaccine design and target site depiction against Ebola viruses: an immunoinformatics study. Scand. J. Immunol. 82, 25–34 (2015).

93. Sadeghi, M. et al. Cytokine expression during early and late phase of acute Puumala hantavirus infection. BMC Immunol. 12, 65 (2011).

94. Nagpal, G. et al. Computer-aided designing of immunosuppressive peptides based on IL-10 inducing potential. Sci. Rep. 7, 42851 (2017).

95. Dhanda, S. K., Vir, P. & Raghava, G. P. S. Designing of interferon-gamma inducing MHC class-II binders. Biol. Direct 8, 30 (2013).

96. Morris, G. M. & Lim-Wilby, M. Molecular Docking BT - Molecular Modeling of Proteins. in (ed. Kukol, A.) 365–382 (Humana Press, 2008). doi:10.1007/978-1-59745-177-2_19

97. Lamiable, A. et al. PEP-FOLD3: faster de novo structure prediction for linear peptides in solution and in complex. Nucleic Acids Res. 44, W449–W454 (2016).

98. Chan, K. F. et al. Divergent T-cell receptor recognition modes of a HLA-I restricted extended tumour-associated peptide. Nat. Commun. 9, 1026 (2018).

99. McMahon, R. M. et al. Structure of HLA-A*0301 in complex with a peptide of proteolipid protein: insights into the role of HLA-A alleles in susceptibility to multiple sclerosis. Acta Crystallogr. D. Biol. Crystallogr. 67, 447–454 (2011).

100. Culshaw, A. et al. Germline bias dictates cross-serotype reactivity in a common dengue-virus-specific CD8+ T cell response. Nat. Immunol. 18, 1228 (2017).

101. Mobbs, J. I. et al. The molecular basis for peptide repertoire selection in the human leucocyte antigen (HLA) C*06:02 molecule. J. Biol. Chem. 292, 17203–17215 (2017).

102. Maenaka, K. et al. Nonstandard Peptide Binding Revealed by Crystal Structures of HLA-B*5101 Complexed with HIV Immunodominant Epitopes. J. Immunol. 165, 3260 LP–3267 (2000).

103. Sun, M. et al. Nα-terminal acetylation for T cell recognition: molecular basis of MHC class I-restricted nα-acetylpeptide presentation. J. Immunol. 192, 5509 LP–5519 (2014).

104. Liu, J. et al. Cross-allele cytotoxic T lymphocyte responses against 2009 pandemic H1N1 influenza A virus among HLA-A24 and HLA-A3 supertype-positive individuals. J. Virol. 86, 13281–13294 (2012).

105. Alpizar, A., Marcilla, M. & Santiago, C. Structure of HLA-B*40:02 in complex with the endogenous peptide REFSKEPEL. TO BE Publ. doi:10.2210/PDB5IEK/PDB

106. Grant, E. J. et al. Broad CD8+ T cell cross-recognition of distinct influenza A strains in humans. Nat. Commun. 9, 5427 (2018).

107. Rosloniec, E. F., Ivey, R. A., Whittington, K. B., Kang, A. H. & Park, H.-W. Crystallographic Structure of a Rheumatoid Arthritis MHC Susceptibility Allele, *HLA-DR1* (*DRB1*0101*), Complexed with the Immunodominant Determinant of Human Type II Collagen. J. Immunol. 177, 3884 LP–3892 (2006).

108. Morris, G. M. et al. AutoDock4 and AutoDockTools4: Automated docking with selective receptor flexibility. J. Comput. Chem. 30, 2785–2791 (2009).

109. Trott, O. & Olson, A. J. AutoDock Vina: improving the speed and accuracy of docking with a new scoring function, efficient optimization, and multithreading. J. Comput. Chem. 31, 455–461 (2010).

110. Tong, J. C., Tan, T. W. & Ranganathan, S. Structure-Based Clustering of Major Histocompatibility Complex (MHC) Proteins for Broad-Based T-Cell Vaccine Design BT - Immunoinformatics. in (eds. De, R. K. & Tomar, N.) 503–511 (Springer New York, 2014). doi:10.1007/978-1-4939-1115-8_27

111. Nezafat, N., Ghasemi, Y., Javadi, G., Khoshnoud, M. J. & Omidinia, E. A novel multi-epitope peptide vaccine against cancer: An in silico approach. J. Theor. Biol. 349, 121–134 (2014).

112. Saha, S. & Raghava, G. P. S. AlgPred: prediction of allergenic proteins and mapping of IgE epitopes. Nucleic Acids Res. 34, W202–W209 (2006).

113. Dimitrov, I., Naneva, L., Doytchinova, I. & Bangov, I. AllergenFP: allergenicity prediction by descriptor fingerprints. Bioinformatics 30, 846–851 (2013).

114. Geourjon, C. & Deléage, G. SOPMA: significant improvements in protein secondary structure prediction by consensus prediction from multiple alignments. Bioinformatics 11, 681–684 (1995).

115. Adiyaman, R. & McGuffin, L. J. Methods for the Refinement of Protein Structure 3D Models. Int. J. Mol. Sci. 20, 2301 (2019).

116. Laskowski, R. A., MacArthur, M. W., Moss, D. S. & Thornton, J. M. PROCHECK: a program to check the stereochemical quality of protein structures. J. Appl. Crystallogr. 26, 283–291 (1993).

117. Laskowski, R. A., Jabłońska, J., Pravda, L., Vařeková, R. S. & Thornton, J. M. PDBsum: Structural summaries of PDB entries. Protein Sci. 27, 129–134 (2018).

118. Wiederstein, M. & Sippl, M. J. ProSA-web: interactive web service for the recognition of errors in three-dimensional structures of proteins. Nucleic Acids Res. 35, W407–W410 (2007).

119. Colovos, C. & Yeates, T. O. Verification of protein structures: patterns of nonbonded atomic interactions. Protein Sci. 2, 1511–1519 (1993).

120. Magnan, C. N., Randall, A. & Baldi, P. SOLpro: accurate sequence-based prediction of protein solubility. Bioinformatics 25, 2200–2207 (2009).

121. Ponomarenko, J. et al. ElliPro: a new structure-based tool for the prediction of antibody epitopes. BMC Bioinformatics 9, 514 (2008).

122. Rodrigues, C. H. M., Pires, D. E. V & Ascher, D. B. DynaMut: predicting the impact of mutations on protein conformation, flexibility and stability. Nucleic Acids Res. 46, W350–W355 (2018).

123. Meng, X.-Y., Zhang, H.-X., Mezei, M. & Cui, M. Molecular docking: a powerful approach for structure-based drug discovery. Curr. Comput. Aided. Drug Des. 7, 146–157 (2011).

124. Yu, H.-T. et al. Hantaan Virus Triggers TLR4-Dependent Innate Immune Responses. Viral Immunol. 25, 387–393 (2012).

125. Kozakov, D. et al. The ClusPro web server for protein–protein docking. Nat. Protoc. 12, 255 (2017).

126. Karplus, M. & McCammon, J. A. Molecular dynamics simulations of biomolecules. Nat. Struct. Biol. 9, 646–652 (2002).

127. Lopéz-Blanco, J. R., Garzón, J. I. & Chacón, P. iMod: multipurpose normal mode analysis in internal coordinates. Bioinformatics 27, 2843–2850 (2011).

128. Tama, F. & Brooks, C. L. SYMMETRY, FORM, AND SHAPE: Guiding Principles for Robustness in Macromolecular Machines. Annu. Rev. Biophys. Biomol. Struct. 35, 115–133 (2006).

129. Meroueh, S. Normal Mode Analysis Theoretical and Applications to Biological and Chemical Systems. Brief. Bioinform. 8, 378–379 (2007).

130. Castiglione, F., Mantile, F., De Berardinis, P. & Prisco, A. How the interval between prime and boost injection affects the immune response in a computational model of the immune system. Comput. Math. Methods Med. 2012, 842329 (2012).

131. Grote, A. et al. JCat: a novel tool to adapt codon usage of a target gene to its potential expression host. Nucleic Acids Res. 33, W526–W531 (2005).

