## Supplementary Tables and Figures for "Computational Approach for Screening the Whole Proteome of Hantavirus and Designing a Multi-Epitope Subunit Vaccine"

### Supplementary Table Legends

**Table S1:** Combined population coverage (%) for the selected T-cell vaccine candidates and their corresponding HLA-alleles.
**Table S2:** Five refined models of the vaccine construct produced by GalaxyRefine sever.
**Table S3:** Scores of the conformational B-cell epitopes of the vaccine construct.
**Table S4:** Lowest energy scores of the 26 vaccine-receptor docked complexes.
**Table S5:** Predicted 70 probable pairs of residues for disulfide engineering with their respective parameters.
**Table S6:** Vibrational entropies for the mutation of the selected pairs of residues for disulfide engineering.

**Table S1:** Combined population coverage (%) for the selected T-cell vaccine candidates and their corresponding HLA-alleles.

| population/area | Class combined | | | population/area | Class combined | | |
|---|---|---|---|---|---|---|---|
| | coverage[a] | average_hit[b] | pc90[c] | | coverage[a] | average_hit[b] | pc90[c] |
| Algeria | 29.62 | 1.12 | 0.14 | Mali | 98.39 | 3.59 | 2.08 |
| Algeria Arab | 29.62 | 1.12 | 0.14 | Mali Black | 98.39 | 3.59 | 2.08 |
| American Samoa | 96.79 | 4.22 | 2.22 | Martinique | 46.01 | 1.02 | 0.19 |
| American Samoa Polynesian | 96.79 | 4.22 | 2.22 | Martinique Black | 46.01 | 1.02 | 0.19 |
| Argentina | 90.56 | 2.71 | 1.02 | Mexico | 99.19 | 4.38 | 2.2 |
| Argentina Amerindian | 89.47 | 2.36 | 0.95 | Mexico Amerindian | 99.48 | 4.37 | 2.34 |
| Argentina Caucasoid | 34.24 | 1.19 | 0.15 | Mexico Mestizo | 90.62 | 3.43 | 1.05 |
| Australia | 97.61 | 4.43 | 1.97 | Mongolia | 80.45 | 2.81 | 0.51 |
| Australia Australian Aborigines | 97.01 | 4.32 | 1.78 | Mongolia Oriental | 80.45 | 2.81 | 0.51 |
| Australia Caucasoid | 98.96 | 4.35 | 2.34 | Morocco | 98.66 | 4.71 | 2.13 |
| Austria | 96.15 | 4.44 | 1.86 | Morocco Arab | 98.66 | 4.67 | 2.12 |
| Austria Caucasoid | 96.15 | 4.44 | 1.86 | Morocco Caucasoid | 98.68 | 4.75 | 2.15 |
| Belgium | 90.23 | 3.39 | 1.02 | Nauru | 0.00 | 0 | 0.4 |
| Belgium Caucasoid | 90.23 | 3.39 | 1.02 | Nauru Micronesian | 0.00 | 0 | 0.4 |
| Bolivia | 36.00 | 1.1 | 0.47 | Netherlands | 29.62 | 0.84 | 0.14 |
| Bolivia Amerindian | 36.00 | 1.1 | 0.47 | Netherlands Caucasoid | 29.62 | 0.84 | 0.14 |
| Borneo | 4.74 | 0.19 | 0.42 | New Caledonia | 99.07 | 5.12 | 3.29 |
| Borneo Austronesian | 4.74 | 0.19 | 0.42 | New Caledonia Melanesian | 99.07 | 5.12 | 3.29 |
| Brazil | 97.19 | 4.16 | 1.78 | New Zealand | 21.92 | 0.67 | 0.13 |
| Brazil Amerindian | 97.39 | 2.86 | 1.45 | New Zealand Polynesian | 21.92 | 0.67 | 0.13 |
| Brazil Caucasoid | 96.99 | 5.01 | 2.17 | Niue | 16.46 | 0.49 | 0.36 |
| Brazil Mixed | 97.67 | 4.53 | 1.98 | Niue Polynesian | 16.46 | 0.49 | 0.36 |
| Brazil Mulatto | 34.55 | 1.05 | 0.15 | North Africa | 97.24 | 4.24 | 1.73 |
| Bulgaria | 98.34 | 4.63 | 2.19 | North America | 98.43 | 5.05 | 2.3 |
| Bulgaria Caucasoid | 93.53 | 3.82 | 1.5 | Northeast Asia | 95.08 | 4.09 | 1.59 |
| Bulgaria Other | 93.00 | 2.83 | 1.14 | Norway | 37.59 | 1.12 | 0.16 |
| Burkina Faso | 76.51 | 1.72 | 0.43 | Norway Caucasoid | 37.59 | 1.12 | 0.16 |

| | | | | | | | |
|---|---|---|---|---|---|---|---|
| Burkina Faso Black | 76.51 | 1.72 | 0.43 | Oceania | 97.22 | 4.56 | 2.19 |
| Cameroon | 97.31 | 3.67 | 1.7 | Oman | 83.57 | 2.51 | 0.61 |
| Cameroon Black | 97.31 | 3.67 | 1.7 | Oman Arab | 83.57 | 2.51 | 0.61 |
| Canada | 2.25 | 0.02 | 0.1 | Pakistan | 88.69 | 2.34 | 0.88 |
| Canada Amerindian | 2.25 | 0.02 | 0.1 | Pakistan Asian | 87.87 | 2.27 | 0.82 |
| Cape Verde | 96.20 | 4.15 | 1.76 | Pakistan Mixed | 90.12 | 2.49 | 1.01 |
| Cape Verde Black | 96.20 | 4.15 | 1.76 | Papua New Guinea | 99.59 | 5.44 | 3.37 |
| Central Africa | 95.99 | 3.52 | 1.46 | Papua New Guinea Melanesian | 99.59 | 5.44 | 3.37 |
| Central African Republic | 66.02 | 2.31 | 0.29 | Paraguay | 0.00 | 0 | 0.7 |
| Central African Republic Black | 66.02 | 2.31 | 0.29 | Paraguay Amerindian | 0.00 | 0 | 0.7 |
| Central America | 19.79 | 0.61 | 0.12 | Peru | 84.73 | 2.79 | 0.66 |
| Chile | 96.50 | 4.26 | 1.76 | Peru Amerindian | 84.42 | 2.77 | 0.64 |
| Chile Amerindian | 99.41 | 5.89 | 3.31 | Peru Mestizo | 1.99 | 0.02 | 0.1 |
| Chile Mixed | 93.60 | 3.41 | 1.25 | Philippines | 96.54 | 4.49 | 2.03 |
| China | 94.95 | 4.05 | 1.55 | Philippines Austronesian | 96.54 | 4.49 | 2.03 |
| China Oriental | 94.95 | 4.05 | 1.55 | Poland | 99.31 | 5.59 | 2.85 |
| Colombia | 15.03 | 0.44 | 0.12 | Poland Caucasoid | 99.31 | 5.59 | 2.85 |
| Colombia Amerindian | 10.79 | 0.31 | 0.11 | Portugal | 97.82 | 5.02 | 2.18 |
| Colombia Black | 22.48 | 0.69 | 0.13 | Portugal Caucasoid | 97.82 | 5.02 | 2.18 |
| Colombia Mestizo | 16.44 | 0.16 | 0.12 | Romania | 93.86 | 3.45 | 1.45 |
| Congo | 20.83 | 0.68 | 0.13 | Romania Caucasoid | 93.86 | 3.45 | 1.45 |
| Congo Black | 20.83 | 0.68 | 0.13 | Russia | 98.98 | 5.55 | 2.74 |
| Cook Islands | 18.96 | 0.57 | 0.12 | Russia Caucasoid | 90.95 | 2.62 | 1.03 |
| Cook Islands Polynesian | 18.96 | 0.57 | 0.12 | Russia Mixed | 75.60 | 1.21 | 0.41 |
| Costa Rica | 11.64 | 0.47 | 0.45 | Russia Other | 99.50 | 6.54 | 3.47 |
| Costa Rica Mestizo | 11.64 | 0.47 | 0.45 | Russia Siberian | 99.15 | 5.66 | 2.92 |
| Croatia | 97.04 | 4.3 | 2.11 | Rwanda | 69.09 | 1.64 | 0.32 |
| Croatia Caucasoid | 97.04 | 4.3 | 2.11 | Rwanda Black | 69.09 | 1.64 | 0.32 |
| Cuba | 96.69 | 5.21 | 2.12 | Samoa | 38.89 | 1.2 | 0.16 |
| Cuba Caucasoid | 91.52 | 3.1 | 1.14 | Samoa Polynesian | 38.89 | 1.2 | 0.16 |

| | | | | | | | |
|---|---|---|---|---|---|---|---|
| Cuba Mixed | 59.04 | 2.16 | 0.24 | Sao Tome and Principe | 91.85 | 3.3 | 1.18 |
| Cuba Mulatto | 92.70 | 2.91 | 1.22 | Sao Tome and Principe Black | 91.85 | 3.3 | 1.18 |
| Czech Republic | 99.27 | 5.47 | 2.75 | Saudi Arabia | 99.15 | 5.28 | 2.34 |
| Czech Republic Caucasoid | 99.30 | 5.51 | 2.78 | Saudi Arabia Arab | 99.15 | 5.28 | 2.34 |
| Czech Republic Other | 34.33 | 1.23 | 0.15 | Scotland | 77.68 | 2.23 | 0.45 |
| Denmark | 24.31 | 0.24 | 0.13 | Scotland Caucasoid | 77.68 | 2.23 | 0.45 |
| Denmark Caucasoid | 24.31 | 0.24 | 0.13 | Senegal | 93.33 | 2.76 | 1.18 |
| East Africa | 94.30 | 3.22 | 1.28 | Senegal Black | 93.33 | 2.76 | 1.18 |
| East Asia | 98.72 | 5.42 | 2.67 | Serbia | 83.03 | 1.88 | 0.59 |
| Ecuador | 81.93 | 3.15 | 0.55 | Serbia Caucasoid | 83.03 | 1.88 | 0.59 |
| Ecuador Amerindian | 81.93 | 3.15 | 0.55 | Singapore | 94.73 | 4.29 | 1.56 |
| England | 98.84 | 5.32 | 2.56 | Singapore Austronesian | 95.01 | 4.32 | 1.54 |
| England Caucasoid | 99.38 | 5.58 | 2.87 | Singapore Oriental | 93.40 | 3.43 | 1.34 |
| England Jew | 44.50 | 0.49 | 0.18 | Slovenia | 40.52 | 1.08 | 0.17 |
| Equatorial Guinea | 16.57 | 0.56 | 0.12 | Slovenia Caucasoid | 40.52 | 1.08 | 0.17 |
| Equatorial Guinea Black | 16.57 | 0.56 | 0.12 | South Africa | 97.63 | 3.47 | 1.66 |
| Ethiopia | 37.59 | 1.5 | 0.64 | South Africa Black | 83.92 | 2.04 | 0.62 |
| Ethiopia Black | 37.59 | 1.5 | 0.64 | South Africa Other | 98.35 | 4.07 | 2.05 |
| Europe | 99.10 | 5.43 | 2.67 | South America | 94.15 | 3.65 | 1.31 |
| Fiji | 15.13 | 0.39 | 0.12 | South Asia | 96.65 | 4.48 | 1.74 |
| Fiji Melanesian | 15.13 | 0.39 | 0.12 | Southeast Asia | 96.26 | 4.59 | 1.96 |
| Finland | 99.62 | 5.24 | 3.12 | Southwest Asia | 93.28 | 3.35 | 1.21 |
| Finland Caucasoid | 99.62 | 5.24 | 3.12 | Spain | 96.35 | 4.08 | 1.61 |
| France | 98.86 | 5.11 | 2.44 | Spain Caucasoid | 96.35 | 4.08 | 1.61 |
| France Caucasoid | 98.86 | 5.11 | 2.44 | Sri Lanka | 23.96 | 0.44 | 0.13 |
| Gabon | 19.36 | 0.69 | 0.12 | Sri Lanka Asian | 23.96 | 0.44 | 0.13 |
| Gabon Black | 19.36 | 0.69 | 0.12 | Sudan | 97.02 | 3.63 | 1.49 |
| Georgia | 99.02 | 5.26 | 2.6 | Sudan Arab | 72.15 | 1.55 | 0.36 |
| Georgia Caucasoid | 99.36 | 5.56 | 2.95 | Sudan Black | 6.49 | 0.13 | 0.21 |
| Georgia Kurd | 98.39 | 4.06 | 2.08 | Sudan Mixed | 91.66 | 3.1 | 1.09 |
| Germany | 99.45 | 5.64 | 2.99 | Sweden | 99.14 | 5.43 | 2.95 |

| | | | | | | | |
|---|---|---|---|---|---|---|---|
| Germany Caucasoid | 99.45 | 5.64 | 2.99 | Sweden Caucasoid | 99.14 | 5.43 | 2.95 |
| Greece | 26.65 | 0.74 | 0.14 | Switzerland | 86.90 | 1.2 | 0.76 |
| Greece Caucasoid | 26.65 | 0.74 | 0.14 | Switzerland Caucasoid | 86.90 | 1.2 | 0.76 |
| Guatemala | 9.60 | 0.16 | 0.11 | Taiwan | 97.80 | 5.13 | 2.56 |
| Guatemala Amerindian | 9.60 | 0.16 | 0.11 | Taiwan Oriental | 97.80 | 5.13 | 2.56 |
| Guinea-Bissau | 89.90 | 2.98 | 0.99 | Thailand | 96.63 | 4.45 | 1.86 |
| Guinea-Bissau Black | 89.90 | 2.98 | 0.99 | Thailand Oriental | 96.63 | 4.45 | 1.86 |
| Hong Kong | 87.91 | 2.94 | 0.83 | Tokelau | 7.84 | 0.24 | 0.33 |
| Hong Kong Oriental | 87.91 | 2.94 | 0.83 | Tokelau Polynesian | 7.84 | 0.24 | 0.33 |
| India | 96.28 | 4.36 | 1.63 | Tonga | 29.44 | 0.88 | 0.43 |
| India Asian | 96.28 | 4.36 | 1.63 | Tonga Polynesian | 29.44 | 0.88 | 0.43 |
| Indonesia | 83.81 | 3.34 | 0.62 | Tunisia | 96.96 | 4.36 | 1.69 |
| Indonesia Austronesian | 83.81 | 3.34 | 0.62 | Tunisia Arab | 96.90 | 4.33 | 1.67 |
| Iran | 98.39 | 4.42 | 2.09 | Tunisia Berber | 38.36 | 1.38 | 0.16 |
| Iran Kurd | 21.68 | 0.74 | 0.13 | Turkey | 58.15 | 1.26 | 0.24 |
| Iran Persian | 98.40 | 4.41 | 2.09 | Turkey Caucasoid | 58.15 | 1.26 | 0.24 |
| Ireland Northern | 99.50 | 5.67 | 2.95 | Uganda | 96.02 | 3.22 | 1.47 |
| Ireland Northern Caucasoid | 99.50 | 5.67 | 2.95 | Uganda Black | 96.02 | 3.22 | 1.47 |
| Ireland South | 99.23 | 5.38 | 2.61 | United Arab Emirates | 13.14 | 0.25 | 0.12 |
| Ireland South Caucasoid | 99.23 | 5.38 | 2.61 | United Arab Emirates Arab | 13.14 | 0.25 | 0.12 |
| Israel | 95.65 | 4.25 | 1.55 | United Kingdom | 84.48 | 1.14 | 0.64 |
| Israel Arab | 98.91 | 5.22 | 2.49 | United Kingdom Caucasoid | 84.48 | 1.14 | 0.64 |
| Israel Jew | 92.55 | 3.83 | 1.19 | United States | 98.45 | 5.08 | 2.32 |
| Italy | 99.33 | 4.92 | 2.67 | United States Amerindian | 98.65 | 4.8 | 2.43 |
| Italy Caucasoid | 99.33 | 4.92 | 2.67 | United States Asian | 98.37 | 5.38 | 2.48 |
| Ivory Coast | 80.37 | 1.74 | 0.51 | United States Austronesian | 20.71 | 0.67 | 0.13 |
| Ivory Coast Black | 80.37 | 1.74 | 0.51 | United States Black | 98.26 | 4.5 | 2.13 |
| Jamaica | 13.14 | 0.53 | 0.46 | United States Caucasoid | 99.33 | 5.57 | 2.83 |

| | | | | | | | |
|---|---|---|---|---|---|---|---|
| Jamaica Black | 13.14 | 0.53 | 0.46 | United States Hispanic | 98.66 | 5 | 2.34 |
| Japan | 98.91 | 5.36 | 2.83 | United States Mestizo | 98.75 | 5.04 | 2.4 |
| Japan Oriental | 98.91 | 5.36 | 2.83 | United States Polynesian | 99.45 | 5.61 | 3.09 |
| Jordan | 85.05 | 2.41 | 0.67 | Venezuela | 92.49 | 3.03 | 1.12 |
| Jordan Arab | 85.05 | 2.41 | 0.67 | Venezuela Amerindian | 92.09 | 3.03 | 1.1 |
| Kenya | 93.97 | 2.71 | 1.21 | Venezuela Caucasoid | 11.45 | 0.12 | 0.11 |
| Kenya Black | 93.97 | 2.71 | 1.21 | Venezuela Mestizo | 9.75 | 0.1 | 0.11 |
| Kiribati | 0.00 | 0 | 0.4 | Vietnam | 94.34 | 4 | 1.44 |
| Kiribati Micronesian | 0.00 | 0 | 0.4 | Vietnam Oriental | 94.34 | 4 | 1.44 |
| Korea; South | 98.50 | 5.28 | 2.44 | Wales | 0.40 | 0 | 0.1 |
| Korea; South Oriental | 98.50 | 5.28 | 2.44 | Wales Caucasoid | 0.40 | 0 | 0.1 |
| Lebanon | 86.59 | 1.77 | 0.75 | West Africa | 96.58 | 3.76 | 1.6 |
| Lebanon Arab | 20.62 | 0.68 | 0.13 | West Indies | 94.61 | 4.02 | 1.52 |
| Lebanon Mixed | 83.11 | 1.09 | 0.59 | World | 97.94 | 4.84 | 2.13 |
| Macedonia | 49.02 | 1.13 | 0.2 | Zambia | 96.18 | 3.32 | 1.63 |
| Macedonia Caucasoid | 49.02 | 1.13 | 0.2 | Zambia Black | 96.18 | 3.32 | 1.63 |
| Malaysia | 78.82 | 2.87 | 0.47 | Zimbabwe | 96.64 | 3.52 | 1.59 |
| Malaysia Austronesian | 51.52 | 1.6 | 0.21 | Zimbabwe Black | 96.64 | 3.52 | 1.59 |
| Malaysia Oriental | 86.79 | 3.66 | 0.76 | | | | |

**Table S2:** Five refined models of the vaccine construct produced by GalaxyRefine sever.

| Model | GDT-HA | RMSD | MolProbity | Clash score | Poor rotamers | Rama favored |
|---|---|---|---|---|---|---|
| Initial | 1.0000 | 0.000 | 3.790 | 179.4 | 4.5 | 88.6 |
| MODEL 1 | 0.9155 | 0.503 | 2.471 | 27.3 | 0.8 | 90.0 |
| MODEL 2 | 0.9228 | 0.503 | 2.553 | 29.8 | 1.2 | 90.2 |
| MODEL 3 | 0.9136 | 0.498 | 2.454 | 27.7 | 0.8 | 90.8 |
| MODEL 4 | 0.9209 | 0.502 | 2.432 | 28.6 | 0.8 | 91.9 |
| MODEL 5 | 0.9289 | 0.490 | 2.514 | 28.6 | 1.2 | 91.0 |

**Table S3:** Scores of the conformational B-cell epitopes of the vaccine construct.

| Serial number | Residues | Number of residues | Score |
|---|---|---|---|
| 1 | _:F41, _:S42, _:I43, _:G44, _:G45, _:G46, _:S47, _:L48, _:G51, _:N52, _:V53, _:L54, _:D55, _:V56, _:G57, _:G58, _:G59, _:S60, _:R61, _:E62, _:Q63, _:K65 | 22 | 0.811 |
| 2 | _:G234, _:P235, _:G236, _:P237, _:G238, _:A239, _:L240, _:L241, _:V242, _:T243, _:F244 | 11 | 0.795 |
| 3 | _:S582, _:D583, _:N584, _:P585, _:C586, _:K587, _:V588, _:D589, _:L590, _:H591, _:K592, _:K593, _:I594, _:L595, _:L596, _:G597, _:S598, _:G604, _:G606, _:Y607, _:F608, _:D609, _:D610, _:L611, _:A612, _:A613, _:K614, _:K615, _:Y616, _:Q617, _:R618, _:T619, _:E620, _:A621, _:D622, _:R623, _:G624, _:F625, _:F626, _:I627, _:T628, _:T632, _:A643, _:W644, _:T645, _:L646, _:K647, _:A648, _:A649, _:A650, _:G651, _:G652, _:G653, _:S654 | 54 | 0.745 |
| 4 | _:E1, _:A2, _:A3, _:A4, _:K5, _:A6, _:P7, _:P8, _:H9, _:A10, _:L11, _:S12, _:E13, _:A14, _:A15, _:A16, _:K17, _:A18, _:K19, _:F20, _:V21, _:A22, _:A23, _:W24, _:T25, _:L26, _:K27, _:A28, _:A29, _:A30, _:G31, _:G32, _:G33, _:S34, _:A35, _:E36, _:L37, _:G38, _:A39, _:F40, _:Q74, _:C75, _:I76, _:Y77, _:T78, _:I79, _:T80, _:S81, _:L82, _:G83, _:G84, _:G85, _:S86, _:M87, _:H106, _:T107, _:A108, _:G109, _:G110, _:G111, _:S112, _:Y113, _:A114, _:Y115, _:S131, _:V132, _:T133, _:L134, _:G135, _:G136, _:G137, _:S138, _:T159, _:F160, _:G161, _:G162, _:G163, _:S164, _:L165, _:P166, _:T167, _:R168, _:R170 | 83 | 0.731 |
| 5 | _:D287, _:G288, _:R289, _:L290, _:N291, _:L292, _:K293, _:G294, _:P295, _:G296, _:P297, _:G298, _:I299, _:L300, _:I301, _:L302, _:S303, _:I304, _:L305, _:L306, _:F307, _:S308, _:F309, _:F310, _:C311, _:P312, _:I313, _:G314, _:P315, _:G316, _:P317, _:G318, _:G319, _:C320, _:Y321, _:R322, _:T323, _:L324, _:N325, _:L326, _:F327, _:R328, _:Y329, _:K330, _:S331, _:R332, _:C333, _:G334, _:P335, _:G336, _:P337, _:G338, _:V339, _:G340, _:L341, _:V342, _:W343, _:G344, _:I345, _:L346, _:L347, _:T348, _:T349, _:E350, _:L351, _:I352, _:I353, _:G354, _:E360, _:A361, _:M447, _:R467, _:L468, _:T469, _:E470, _:V473, _:K483, _:P485, _:V486, _:P487, _:L488, _:G489, _:Q490, _:V491, _:T492, _:D493, _:L494, _:K495, _:I496, _:E497, _:S498, _:S499, _:C500, _:N501, _:F502, _:D503, _:K504, _:K505, _:R506, _:I507, _:E508, _:W509, _:K510, _:D511, _:P512, _:D513, _:G514, _:M515, _:L516, _:R517 | 110 | 0.724 |
| 6 | _:C245, _:F246, _:G247, _:W248 | 4 | 0.713 |
| 7 | _:D193, _:G194, _:P195, _:G196, _:P197, _:G198, _:P199, _:I200, _:Y206, _:M207, _:L208, _:S209, _:T210, _:R211, _:G212, _:G214, _:P215, _:G216, _:P217, _:G218, _:Q219, _:R220 | 22 | 0.704 |
| 8 | _:M188, _:G189, _:I190, _:Q191, _:L192 | 5 | 0.689 |
| 9 | _:T67, _:Q68, _:M69, _:G70, _:G71, _:G72 | 6 | 0.609 |
| 10 | _:L406, _:K407, _:K408, _:K409, _:S410, _:S411, _:Y413, _:S415, _:T546, _:C547, _:K548, _:K550, _:H551 | 13 | 0.557 |

**Table S4:** Lowest energy scores of the 26 vaccine-receptor docked complexes.

| Cluster | Members | Representative | Weighted Score |
|---|---|---|---|
| 0 | 55 | Center | -1113.8 |
| 0 | 55 | Lowest Energy | -1137.6 |
| 1 | 35 | Center | -1000.8 |
| 1 | 35 | Lowest Energy | -1246.6 |
| 2 | 29 | Center | -1074.2 |
| 2 | 29 | Lowest Energy | -1215.8 |
| 3 | 23 | Center | -1029.7 |
| 3 | 23 | Lowest Energy | -1104.8 |
| 4 | 21 | Center | -964.9 |
| 4 | 21 | Lowest Energy | -1078 |
| 5 | 21 | Center | -1197.2 |
| 5 | 21 | Lowest Energy | -1197.2 |
| 6 | 19 | Center | -988.8 |
| 6 | 19 | Lowest Energy | -1112.1 |
| 7 | 17 | Center | -1135.2 |
| 7 | 17 | Lowest Energy | -1135.2 |
| 8 | 17 | Center | -1010.3 |
| 8 | 17 | Lowest Energy | -1132.6 |
| 9 | 17 | Center | -967.2 |
| 9 | 17 | Lowest Energy | -1104.1 |
| 10 | 15 | Center | -1038.9 |
| 10 | 15 | Lowest Energy | -1110.1 |
| 11 | 15 | Center | -1136.3 |
| 11 | 15 | Lowest Energy | -1292 |
| 12 | 14 | Center | -1124.7 |
| 12 | 14 | Lowest Energy | -1125.6 |
| 13 | 14 | Center | -1029.7 |
| 13 | 14 | Lowest Energy | -1234.3 |
| 14 | 13 | Center | -1135.9 |
| 14 | 13 | Lowest Energy | -1158.4 |
| 15 | 13 | Center | -982.3 |
| 15 | 13 | Lowest Energy | -1098.5 |
| 16 | 12 | Center | -1005.7 |
| 16 | 12 | Lowest Energy | -1048.5 |
| 17 | 12 | Center | -1055.6 |
| 17 | 12 | Lowest Energy | -1085.6 |
| 18 | 11 | Center | -1121.6 |
| 18 | 11 | Lowest Energy | -1121.6 |

| | | | |
|---|---|---|---|
| 19 | 10 | Center | -1154.2 |
| 19 | 10 | Lowest Energy | -1154.2 |
| 20 | 10 | Center | -1039.4 |
| 20 | 10 | Lowest Energy | -1039.4 |
| 21 | 9 | Center | -990 |
| 21 | 9 | Lowest Energy | -1112.8 |
| 22 | 7 | Center | -968.5 |
| 22 | 7 | Lowest Energy | -1079.6 |
| 23 | 7 | Center | -984.2 |
| 23 | 7 | Lowest Energy | -991.4 |
| 24 | 5 | Center | -1052.6 |
| 24 | 5 | Lowest Energy | -1052.6 |
| 25 | 4 | Center | -1071.2 |
| 25 | 4 | Lowest Energy | -1071.2 |

**Table S5:** Predicted 70 probable pairs of residues for disulfide engineering with their respective parameters.

| Res1 Seq # | Res1 AA | Res2 Seq # | Res2 AA | Chi3 | Energy | Sum B-Factors |
|---|---|---|---|---|---|---|
| 344 | GLY | 507 | ILE | -81.14 | 0.62 | 0 |
| 184 | ARG | 265 | PRO | 86.53 | 0.84 | 0 |
| 24 | TRP | 87 | MET | -95.48 | 1.08 | 0 |
| 401 | THR | 436 | GLY | 80.96 | 1.47 | 0 |
| 333 | CYS | 500 | CYS | 99.62 | 1.6 | 0 |
| 413 | TYR | 556 | GLU | 80.88 | 1.72 | 0 |
| 426 | ASN | 570 | LYS | -109.84 | 1.77 | 0 |
| 378 | GLY | 422 | VAL | 110.52 | 1.84 | 0 |
| 270 | ALA | 375 | PRO | -79.7 | 1.88 | 0 |
| 373 | ARG | 379 | ALA | 106.34 | 1.9 | 0 |
| 94 | PHE | 97 | GLY | 112.83 | 2.16 | 0 |
| 434 | VAL | 557 | SER | 112.98 | 2.29 | 0 |
| 384 | PHE | 393 | ARG | 103.31 | 2.39 | 0 |
| 88 | PRO | 157 | LEU | 96.32 | 2.52 | 0 |
| 109 | GLY | 165 | LEU | 116.72 | 2.6 | 0 |
| 429 | SER | 566 | CYS | -99.83 | 2.62 | 0 |
| 616 | TYR | 621 | ALA | 75.84 | 2.74 | 0 |
| 486 | VAL | 489 | GLY | 100.1 | 2.8 | 0 |
| 374 | GLY | 398 | THR | 68.68 | 2.99 | 0 |
| 294 | GLY | 507 | ILE | 67.01 | 3.07 | 0 |
| 374 | GLY | 562 | ARG | -63.23 | 3.12 | 0 |
| 644 | TRP | 653 | GLY | 107.6 | 3.14 | 0 |
| 112 | SER | 115 | TYR | 107.52 | 3.2 | 0 |
| 320 | CYS | 348 | THR | 93.24 | 3.21 | 0 |
| 47 | SER | 51 | GLY | -81.94 | 3.3 | 0 |
| 284 | HIS | 360 | GLU | -108.84 | 3.48 | 0 |
| 597 | GLY | 606 | GLY | 120.9 | 3.49 | 0 |
| 309 | PHE | 345 | ILE | -86.99 | 3.61 | 0 |
| 315 | PRO | 319 | GLY | -77.41 | 3.67 | 0 |
| 613 | ALA | 646 | LEU | 125.62 | 3.73 | 0 |
| 544 | GLU | 558 | TYR | 99.29 | 3.88 | 0 |
| 95 | LEU | 149 | GLY | -110.57 | 4 | 0 |
| 204 | ALA | 262 | CYS | -68.07 | 4.08 | 0 |
| 86 | SER | 158 | SER | 85.16 | 4.16 | 0 |
| 104 | PRO | 170 | ARG | 111.31 | 4.18 | 0 |
| 29 | ALA | 33 | GLY | 118.09 | 4.39 | 0 |
| 144 | PHE | 225 | TRP | 82.01 | 4.39 | 0 |

| | | | | | | |
|---|---|---|---|---|---|---|
| 397 | LYS | 420 | ALA | -114.22 | 4.5 | 0 |
| 377 | PRO | 422 | VAL | 84.84 | 4.62 | 0 |
| 454 | GLN | 521 | ASN | -105.37 | 4.63 | 0 |
| 282 | LEU | 364 | GLY | -97.76 | 4.71 | 0 |
| 362 | ASP | 447 | MET | -86.3 | 4.71 | 0 |
| 285 | TRP | 361 | ALA | -93.09 | 4.72 | 0 |
| 524 | VAL | 536 | LYS | -88.78 | 4.78 | 0 |
| 279 | VAL | 365 | PHE | -100.67 | 4.8 | 0 |
| 180 | SER | 570 | LYS | 96.64 | 4.84 | 0 |
| 178 | GLY | 572 | LEU | 90.07 | 5.12 | 0 |
| 487 | PRO | 511 | ASP | 119.02 | 5.2 | 0 |
| 148 | GLY | 256 | GLY | 118.64 | 5.22 | 0 |
| 618 | ARG | 621 | ALA | -79.61 | 5.25 | 0 |
| 467 | ARG | 487 | PRO | 106.53 | 5.34 | 0 |
| 615 | LYS | 622 | ASP | 96.78 | 5.38 | 0 |
| 309 | PHE | 347 | LEU | 86.08 | 5.43 | 0 |
| 75 | CYS | 171 | LEU | 120.41 | 5.5 | 0 |
| 146 | TYR | 258 | GLY | -110.65 | 5.68 | 0 |
| 376 | GLY | 431 | VAL | 93.07 | 5.85 | 0 |
| 374 | GLY | 431 | VAL | -99.59 | 5.94 | 0 |
| 77 | TYR | 170 | ARG | -76.54 | 5.99 | 0 |
| 435 | ILE | 557 | SER | -90.58 | 6.54 | 0 |
| 210 | THR | 218 | GLY | -102.85 | 6.55 | 0 |
| 271 | LEU | 381 | ARG | 120.18 | 6.67 | 0 |
| 286 | PHE | 351 | LEU | -112.5 | 6.78 | 0 |
| 488 | LEU | 511 | ASP | -59.65 | 6.79 | 0 |
| 475 | PHE | 482 | LYS | 94.32 | 6.89 | 0 |
| 48 | LEU | 51 | GLY | 70.91 | 6.98 | 0 |
| 443 | GLY | 458 | MET | -57.84 | 7.01 | 0 |
| 278 | GLY | 368 | THR | -62.57 | 7.79 | 0 |
| 228 | MET | 262 | CYS | -59.8 | 7.92 | 0 |
| 445 | ASN | 456 | LEU | -75.3 | 8.64 | 0 |
| 283 | GLY | 475 | PHE | 124.5 | 9.1 | 0 |

**Table S6:** Vibrational entropies for the mutation of the selected pairs of residues for disulfide engineering.

| Res1 Seq # | Res1 AA | ΔΔG DynaMut (kcal/mol) | Res2 Seq # | Res2 AA | ΔΔG DynaMut (kcal/mol) | Chi3 | Energy | Sum B-Factors |
|---|---|---|---|---|---|---|---|---|
| 344 | GLY | 2.106 | 507 | ILE | -1.297 | -81.14 | 0.62 | 0 |
| 184 | ARG | -0.411 | 265 | PRO | -0.018 | 86.53 | 0.84 | 0 |
| 401 | THR | 1.001 | 436 | GLY | 1.571 | 80.96 | 1.47 | 0 |
| 413 | TYR | -0.499 | 556 | GLU | -0.814 | 80.88 | 1.72 | 0 |
| 270 | ALA | -0.082 | 375 | PRO | -0.402 | -79.7 | 1.88 | 0 |

### **Supplementary Figure Legends**

**Figure S1:** *In silico* simulation of immune response after injecting three injections by 4-weeks apart. (a) PLB cell population. (b) TR (regulatory) cell population per state. (c) NK cell population. (d) EP population per state.

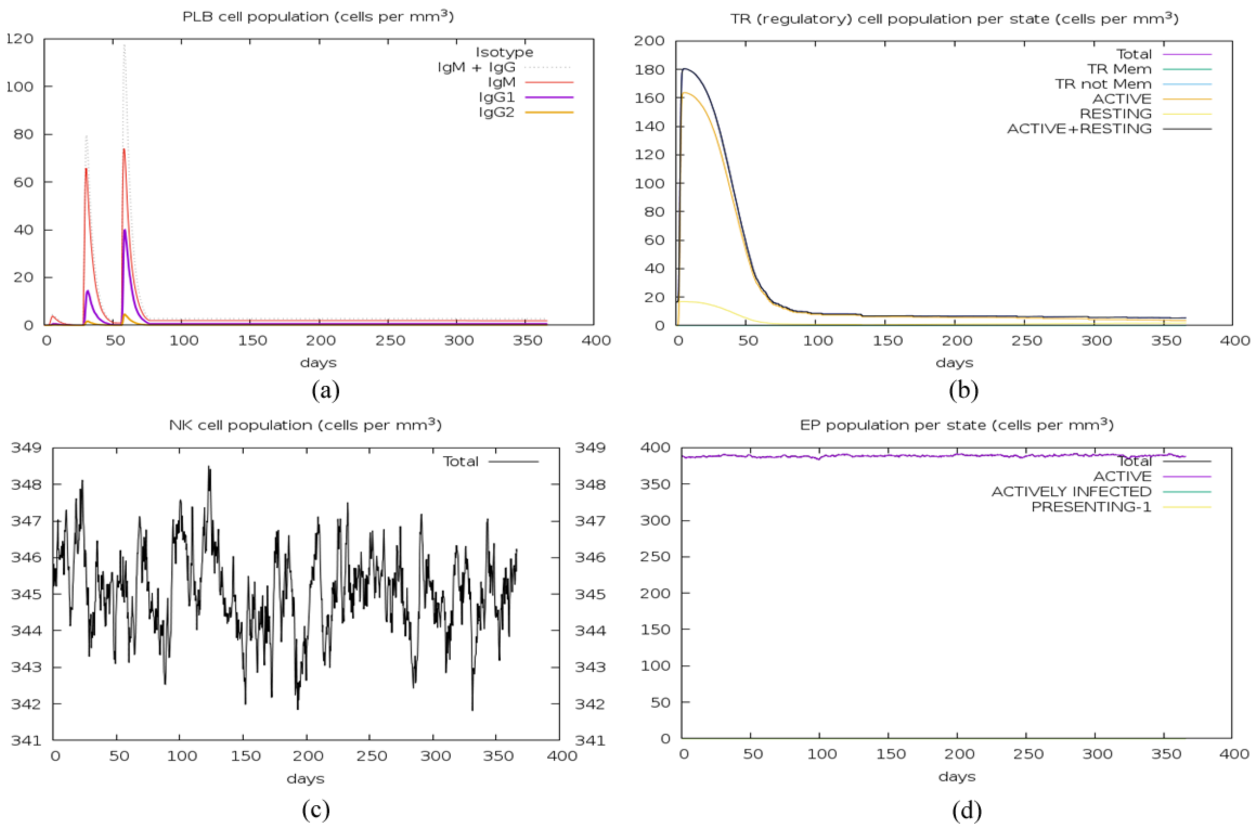


**Figure S1:** *In silico* simulation of immune response after injecting three injections by 4-weeks apart. (a) PLB cell population. (b) TR (regulatory) cell population per state. (c) NK cell population. (d) EP population per state.
